# Brain gene co-expression networks link complement signaling with convergent synaptic pathology in schizophrenia

**DOI:** 10.1101/2020.03.03.975722

**Authors:** Minsoo Kim, Jillian R. Haney, Pan Zhang, Leanna M. Hernandez, Lee-kai Wang, Laura Perez-Cano, Loes M. Olde Loohuis, Luis de la Torre-Ubieta, Michael J. Gandal

**Affiliations:** Department of Psychiatry, David Geffen School of Medicine, University of California, Los Angeles, Los Angeles, CA, USA; Department of Human Genetics, David Geffen School of Medicine, University of California, Los Angeles, Los Angeles, CA, USA; Program in Neurobehavioral Genetics, Semel Institute, David Geffen School of Medicine, University of California, Los Angeles, Los Angeles, CA, USA; Department of Neurology, David Geffen School of Medicine, University of California, Los Angeles, Los Angeles, CA, USA; STALICLA DDS, Barcelona, Spain

## Abstract

The most significant common variant association for schizophrenia (SCZ) reflects increased expression of the complement component 4A (*C4A*). Yet, it remains unclear how *C4A* interacts with other SCZ risk genes and whether the complement system is more broadly implicated in SCZ pathogenesis. Here, we integrate several existing, large-scale genetic and transcriptomic datasets to interrogate the functional role of the complement system and *C4A* in the human brain. Surprisingly, we find no significant genetic enrichment among known complement system genes for SCZ. Conversely, brain co-expression network analyses using *C4A* as a seed gene revealed that genes down-regulated when *C4A* expression increased exhibit strong and specific genetic enrichment for SCZ risk. This convergent genomic signal reflected neuronal, synaptic processes and was sexually dimorphic and most prominent in frontal cortical brain regions. Overall, these results indicate that synaptic pathways—rather than the complement system—are the driving force conferring SCZ risk.

## Introduction

SCZ is a highly heritable and disabling neurodevelopmental, psychiatric disorder that affects ∼1% of the general population^1,2^. Despite tremendous contribution to public health burden worldwide, there have been no fundamental advances in the treatment of SCZ since the 1980s, due in large part to the lack of novel, robust therapeutic targets. The recent success of genome-wide association studies (GWAS)^3-5^ brings hope that genetics can provide novel insights into underlying disease mechanisms and identify new biological pathways for intervention. However, the transition from GWAS to mechanistic insights is challenged by daunting levels of polygenicity and small effect sizes of associated variants^2,6^. One potential solution has been to incorporate GWAS results within the context of known molecular and cellular pathways, leveraging prior knowledge that genes do not act in isolation, to identify biological processes exhibiting robust genetic convergence^7,8^.

The strongest and first-identified GWAS signal for SCZ lies in the major histocompatibility complex (MHC) region, traditionally known for its role in immunity. This association was subsequently shown to reflect in part complex genetic variation within the *C4* locus^9^, where human *C4* is encoded by two genes—*C4A* and *C4B*—which exist in different combinations of copy numbers, commonly ranging from zero to four copies of each gene per individual. Previous work demonstrated that such multiallelic copy number variation (mCNV) of *C4* influences gene expression and that elevated expression of *C4A*, but not *C4B*, confers SCZ risk^9^. *C4A* encodes an early component of the classical complement pathway, a part of the innate immune system that serves to clear cellular debris and provide the first line of antimicrobial defense. The strength and novelty of this association has prompted speculation that *C4A*—and the complement system more broadly—may represent a novel therapeutic target for SCZ. However, apart from *C4A*, surprisingly little is known about the broader relevance of the complement system in SCZ pathogenesis. Furthermore, it remains unclear whether *C4A* interacts with other established risk factors.

Within the brain, the complement system plays a distinct, non-inflammatory role as a mediator of synaptic pruning^9-11^, where it tags synapses for microglia-dependent elimination. Intriguingly, excessive pruning has long been hypothesized in SCZ^12-14^ and thought to reflect reduced cortical thickness^15^ as well as dendritic spine abnormalities^16^ observed in SCZ cases. However, these links have yet to be proven or tied to a concrete genetic mechanism. Complicating matters, the lack of evolutionary conservation of *C4A* has hindered direct investigation in model organisms. Whereas human stem cell-based assays have been used to study aspects of synapse elimination relevant to SCZ^17^, these systems fail to recapitulate the complete range of neuronal-glial interactions present in the human brain, nor have they been shown to reach postnatal levels of maturity^18^ when pruning largely occurs. As such, we reasoned that direct assessment in the human brain is an important first step to elucidate the specific molecular processes through which *C4A* increases risk for disease.

In this study, we integrated large-scale genetic and brain transcriptomic datasets from PsychENCODE^19,20^ and GTEx^21^ to interrogate the functional role of *C4A* in the human brain and its relation to other SCZ risk factors. We used gene co-expression networks to capture coherent biological processes that covary across samples^22^ and hence provide an unbiased functional annotation for *C4A*. We took a seeded approach, identifying genes whose expression is either positively or negatively correlated with *C4A* expression. Genes positively correlated with *C4A* captured the known complement components as well as astrocyte, microglial, and NFkB signaling pathways, but they showed no genetic enrichment for SCZ. In contrast, genes negatively correlated with *C4A* reflected neuronal and synaptic pathways and exhibited strong convergent enrichment for SCZ genetic risk. Altogether, these results highlight the human brain-specific function of *C4A* and provide evidence for complex interplay between *C4A* and synaptic processes to confer SCZ risk.

## Results

### Limited evidence for SCZ genetic association within the known complement system

We first sought to determine whether genetic evidence supported SCZ association for any of the 57 genes annotated within the complement system (Methods). As GWAS loci are difficult to definitively map to causal genes, we assessed several lines of evidence supporting a putative association (**Fig. 1a** and **Supplementary Table 1**). We first evaluated the proximity of these genes to SCZ GWAS loci^4,5^. Outside the MHC region, nine genes were within 1 Mb of genome-wide significant loci (**Fig. 1a**). Of these, three were not considered brain-expressed in PsychENCODE^19^, and several were within the same genomic region. Three genes—*CD46, CSMD1*, and *CLU*—were the closest gene to their respective index single-nucleotide polymorphism (SNP). *CSMD1* and *CD46* had support from Hi-C interactions in fetal and adult brain^23^, and *CLU* and *CD46* had additional support from summary-data-based Mendelian randomization (SMR)^24^ at FDR < 0.05 and *P*_HEIDI_ > 0.05 (**Fig. 1a** and **Supplementary Table 1**). Altogether, these findings provided a moderate level of evidence supporting SCZ association for up to four genes within the complement system.

**Fig. 1.**
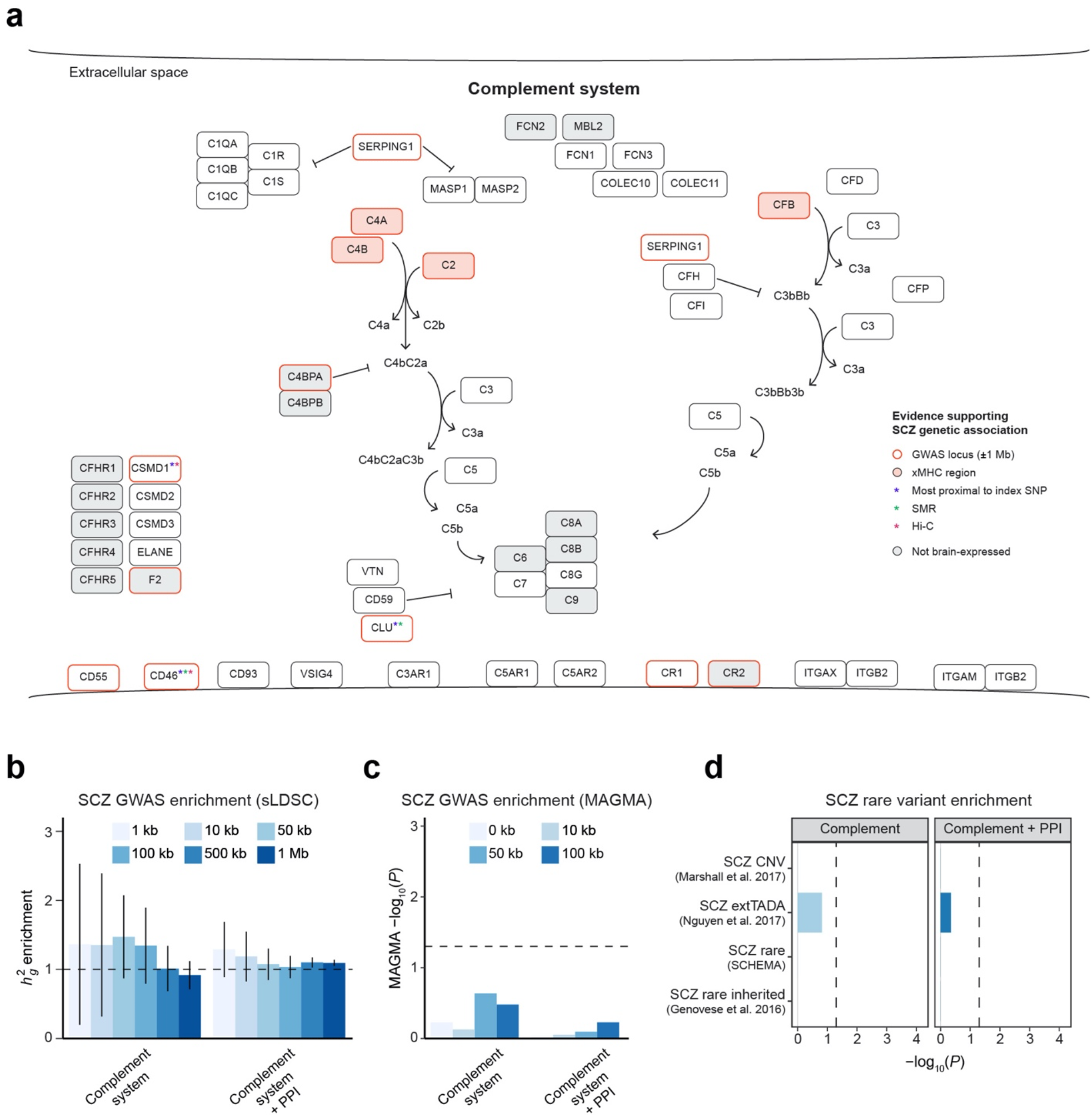
Limited evidence for broad genetic enrichment within the complement system. **a**, The complement system is composed of 57 genes which function together in a cascade to clear cellular debris, opsonize microbes, and mediate synaptic pruning. Here, we plot genes annotated within the complement system and corresponding evidence for SCZ genetic association^4,5^, based on proximity to GWAS loci, support from SMR (summary-data-based Mendelian randomization), and Hi-C interactions in fetal and adult brain. No enrichment of SCZ GWAS signals was observed for the complement system or an expanded annotation including high-confidence PPIs (InWeb3; n = 545 genes), using **b**, sLDSC or **c**, MAGMA with varying window sizes around each gene. All error bars denote standard errors of estimates of heritability enrichment, where the enrichment is defined as the proportion of SNP-based heritability over proportion of SNPs. **d**, The complement system did not show enrichment for SCZ risk genes from rare variant studies.

To determine whether this putative association of four complement system genes is greater than expected by chance, and to test whether the complement system as a whole is broadly enriched for SCZ GWAS signals, we used stratified LD score regression (sLDSC)^25^. We found no significant enrichment of SNP-based heritability in SCZ, testing a range of window sizes around each gene (**Fig. 1b**). A similar lack of enrichment was found using a second method, MAGMA^26^ (**Fig. 1c**). To account for the small number of genes in this pathway, we further expanded the annotation to include high-confidence protein-protein interactions (PPIs)^27^ for the complement system and still observed no significant enrichment. Finally, we tested whether any of these gene sets were enriched for genes implicated in SCZ through rare variant association studies, again finding no evidence of enrichment (**Fig. 1d**). These included genes within the eight recurrent CNV regions associated with SCZ^28^ and genes harboring an excess of rare, likely gene-disrupting (LGD) variants in SCZ probands^29,30^. Together, these results do not support broad genetic association for SCZ within the complement system.

### Seeded co-expression networks provide brain-specific functional annotation for *C4A*

The previous analyses relied on known gene set annotations which are often incomplete, especially for biological processes occurring in the human brain^31^. Additionally, the non-inflammatory role of *C4A*—and the complement system—as an effector of synaptic pruning may not be fully reflected in these annotations. To address this, we turned to gene co-expression network analyses, which can provide an orthogonal, unbiased functional annotation based on correlated gene expression patterns across samples^22^. Here, we took a ‘seeded’ approach, identifying genes either positively or negatively correlated with *C4A* expression and using such ‘guilt-by-association’ to draw biological inference.

We first constructed a *C4A*-seeded co-expression network from frontal cortex samples of neurotypical controls in PsychENCODE^19,20^ (**Fig. 2a**). To mitigate the potential influence of germline mCNV, we imputed *C4* structural alleles from nearby SNP genotypes^9^ in individuals of European ancestry (N = 812; **Extended Data Fig. 1**). We then selected control samples with high-quality imputation results carrying the most common diploid *C4A* copy number (CN = 2, N = 145; **Extended Data Fig. 2**; Methods). Using these samples, we identified 3,021 genes co-expressed with *C4A* at FDR < 0.05 (**Supplementary Table 2**). These included 1,869 positively co-expressed genes as well as 1,152 negatively co-expressed genes (herein referred to as “*C4A*-positive” and “*C4A*-negative” genes). As a positive control, the known complement signaling pathway was overrepresented among *C4A*-positive genes (odds ratio (OR) = 17.2, *P* < 10^−16^), but not *C4A*-negative genes (OR = 0, *P* = 1). In addition, *C4A*-positive genes were most strongly enriched for “immune effector process” and “response to cytokine” Gene Ontology (GO) terms (FDR’s < 10^−41^), whereas *C4A*-negative genes were most strongly enriched for “anterograde trans-synaptic signaling” and “chemical synaptic transmission” GO terms (FDR’s < 10^−12^).

**Fig. 2.**
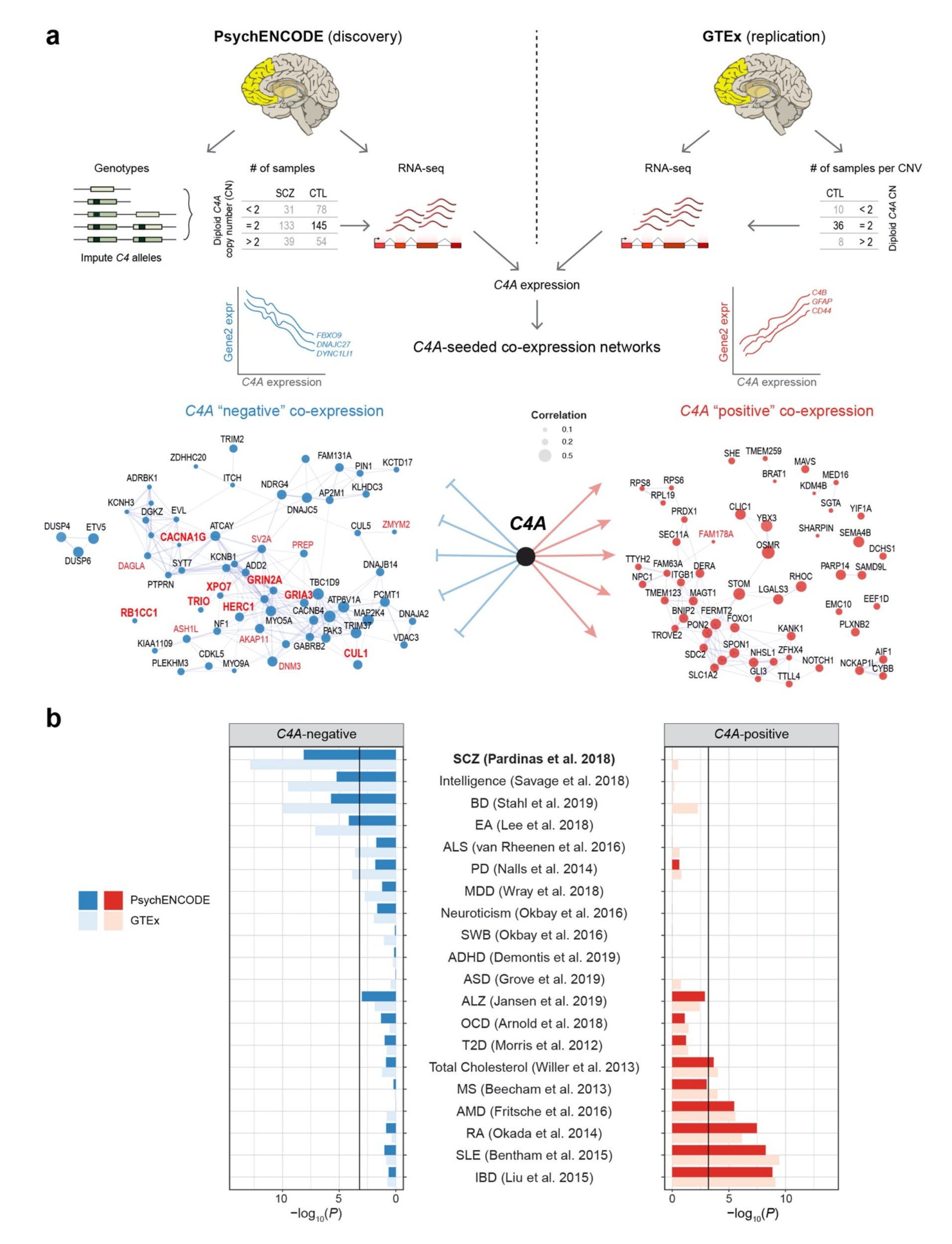
*C4A*-seeded co-expression networks capture convergent genetic risk for SCZ. **a**, Overview of the generation of *C4A*-seeded networks, using control samples from PsychENCODE and GTEx. Node size is proportional to |correlation| with *C4A* expression and edges represent gene-gene co-expression. Shown in red labels are SCZ risk genes from SCHEMA^29^ reaching FDR or exome-wide (bold) significance. **b**, *C4A*-positive and *C4A*-negative genes showed enrichment for distinct GWAS signals, where *C4A*-negative, but not *C4A*-positive, genes showed enrichment for SNP-based heritability in SCZ. Results replicated in the independent GTEx dataset. The black line denotes Bonferroni-adjusted *P* value at 0.05/80. ADHD (attention-deficit/hyperactivity disorder), ALS (amyotrophic lateral sclerosis), ALZ (Alzheimer disease), AMD (age-related macular degeneration), ASD (autism spectrum disorder), BD (bipolar disorder), EA (educational attainment), IBD (inflammatory bowel disease), MDD (major depressive disorder), MS (multiple sclerosis), OCD (obsessive-compulsive disorder), PD (Parkinson’s disease), RA (rheumatoid arthritis), SLE (systemic lupus erythematosus), SWB (subjective well-being), T2D (type 2 diabetes).

For replication, we generated an analogous seeded network in the independent GTEx dataset^21^. We observed highly significant overlap among *C4A*-positive and *C4A*-negative genes across these datasets (OR’s = 19 and 16, *P*’s < 10^−16^, respectively; **Extended Data Fig. 3**). As an additional control, we generated 10,000 seeded networks using randomly sampled seed genes (Methods). The original *C4A*-positive network showed greater enrichment for the known complement components than 98% of all other networks generated in this manner (**Extended Data Fig. 4a**).

### *C4A*-negative, but not *C4A*-positive, genes show strong SCZ genetic enrichment

We next sought to determine whether this network-based, brain-specific functional annotation for *C4A* better captured convergent genetic risk for SCZ. Consistent with our results above, we did not find enrichment of SNP-based heritability for SCZ among *C4A*-positive genes (**Fig. 2b**). These genes were instead associated with autoimmune and chronic inflammatory conditions, such as inflammatory bowel disease (IBD), rheumatoid arthritis (RA), and lupus (SLE). In contrast, *C4A*-negative genes were strongly enriched for SNP-based heritability in SCZ and in several other neuropsychiatric disorders, to a lesser degree (**Fig. 2b**). These findings were replicated in GTEx, so we subsequently combined both networks from PsychENCODE and GTEx to yield a high-confidence seeded network (Methods). Notably, in this network, among the ten genes harboring rare loss-of-function variants in SCZ probands at exome-wide significance^29^, eight were negatively co-expressed with *C4A* at FDR < 0.1 (*TRIO, GRIN2A, XPO7, CUL1, GRIA3, HERC1, RB1CC1, CACNA1G*), suggesting convergence of polygenic effects across the allelic spectrum (logistic regression, FDR = 9.0 × 10^−4^; **Fig. 2a** and **Extended Data Fig. 4b**; Methods). The remaining two genes (*SETD1A, SP4*) show peak expression in the fetal brain, suggesting alternative developmental mechanisms^32^.

### Network expansion with increased *C4A* copy number

*C4A* expression is likely influenced by both genetic and environmental factors. In PsychENCODE, we observed that ∼22% of the variation in *C4A* expression can be explained by germline mCNV (**Extended Data Fig. 5**). However, it remains unknown what effect these genetic factors have on *C4A* co-expression. To address this, we stratified all PsychENCODE samples with high-quality imputation results (N = 552) into three CNV groups based on diploid *C4A* copy number of < 2, 2, and > 2, representing a gradient of increasing genetic risk for SCZ (**Extended Data Fig. 2**; Methods). We then generated *C4A*-seeded networks for each group, using bootstrap to match the sample size (100 samples + 10,000 iterations). Remarkably, we observed a dramatic increase in network size as *C4A* copy number increased (**Fig. 3a**). With increased genomic copy number, the number of both *C4A*-positive and *C4A*-negative genes was substantially larger, indicating that *C4A* is more strongly connected and likely plays more of a driver role (**Extended Data Fig. 6**). This network expansion was preserved across a range of correlation and FDR thresholds (**Fig. 3b**) and was not associated with technical factors such as postmortem interval (PMI) or RIN. Furthermore, this network expansion was not observed for *C4B*-seeded networks, demonstrating the specificity of this association (**Fig. 3c** and **Extended Data Fig. 7**; Methods). Together, these results show that genotypes conferring increased risk for SCZ are associated with distinct brain gene co-expression networks.

**Fig. 3.**
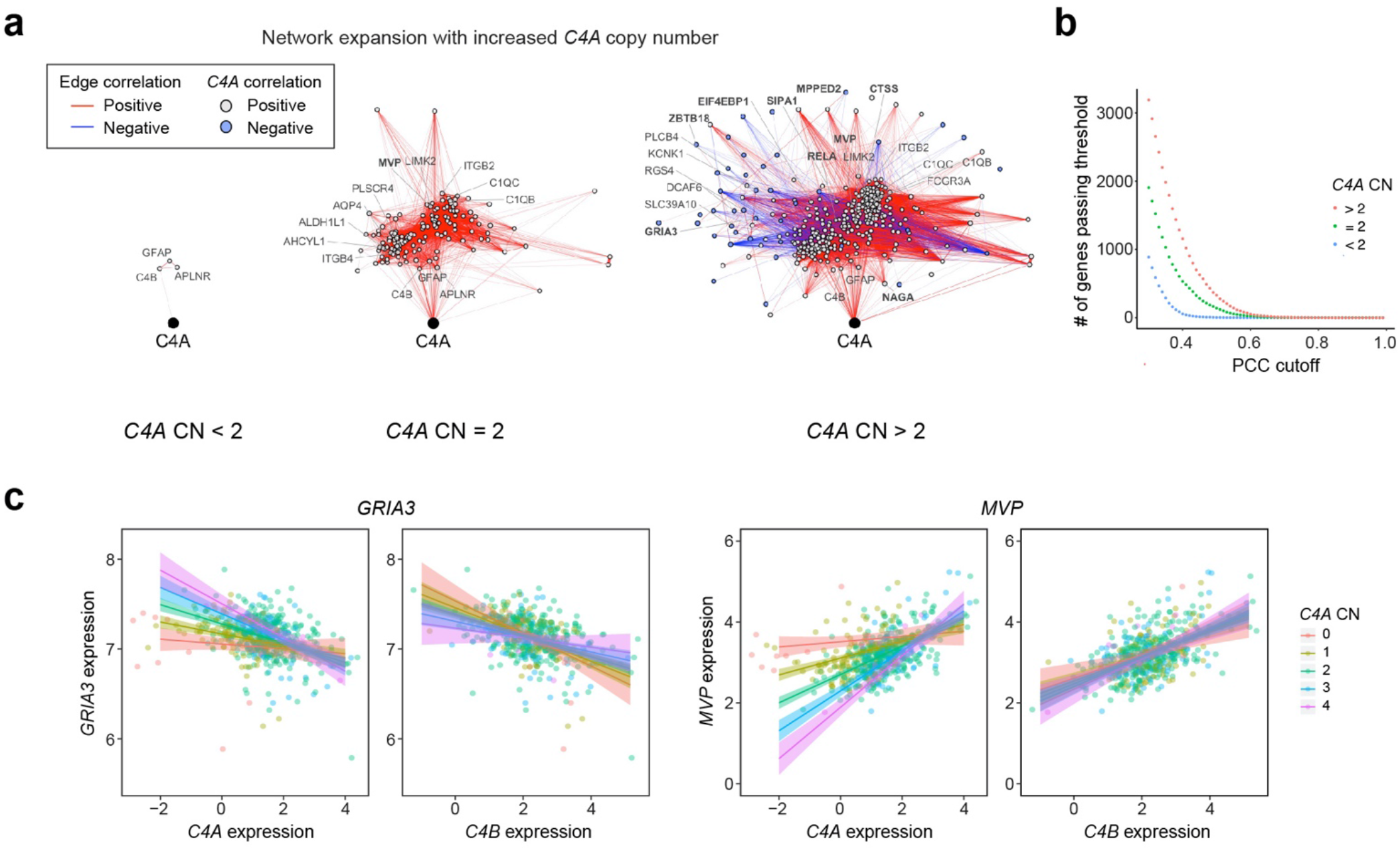
Strong network expansion with increased *C4A* copy number. **a**, *C4A*-seeded co-expression networks were generated following stratification of the PsychENCODE dataset by imputed *C4A* copy number. A substantial network expansion was observed with increased *C4A* copy number. Each network was generated via bootstrap (100 samples, 10,000 iterations) for robustness. Edges represent Pearson’s correlation coefficient (PCC) > 0.5 and edge weights represent the strength of the correlation. Probable SCZ risk genes implicated by common or rare variant studies are highlighted in bold. **b**, *C4A*-seeded networks expanded in size regardless of the applied PCC threshold. **c**, The nonlinear network expansion was specific to *C4A* as a seed gene, and not observed for *C4B*. Two genes, *GRIA3* and *MVP*, are shown to illustrate this specificity. Shown are fitted linear models with 95% confidence bands.

### Seeded networks capture *C4A*-associated pathways and cell-types

We then sought to understand the biological pathways and cell-types captured by these *C4A*-seeded networks. As above, *C4A*-positive and *C4A*-negative genes were enriched for distinct GO terms: *C4A*-positive genes for inflammatory pathways and *C4A*-negative genes for synapse-related pathways (**Supplementary Fig. 1**). Overlap of these genes with a set of previously characterized brain co-expression modules^19^ confirmed their broad relationship to inflammatory and synaptic function, respectively (**Fig. 4a** and **Supplementary Fig. 2**).

**Fig. 4.**
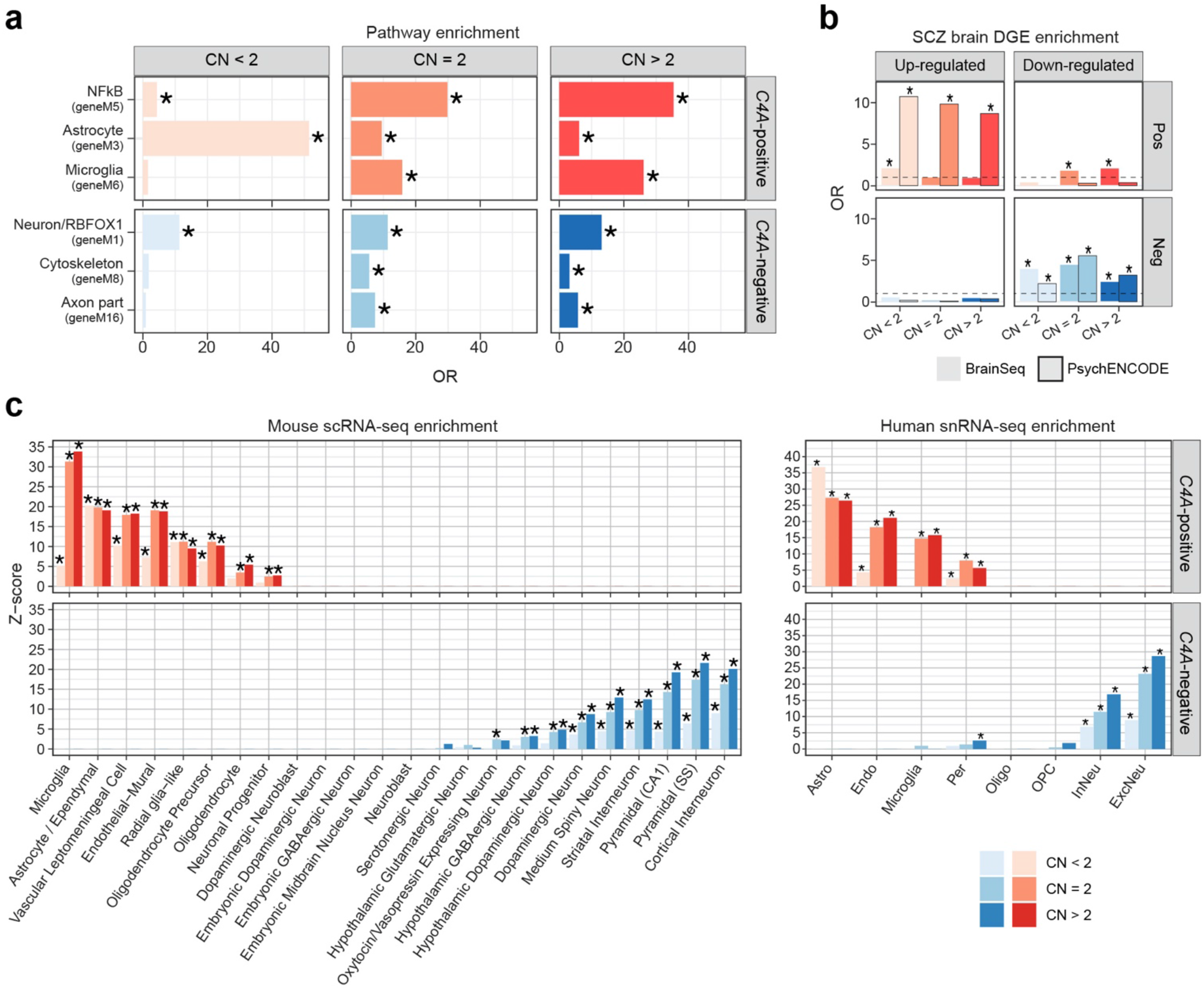
*C4A*-seeded co-expression networks identify transcriptional correlates of synaptic pruning. **a**, The top *C4A*-positive and *C4A*-negative genes showed distinct enrichments for neurobiological pathways and cell-types. With increasing *C4A* copy number, *C4A*-positive genes showed greater enrichment for microglia and NFkB pathways, while *C4A*-negative genes showed greater enrichment for neuron- and synapse-related modules. OR = odds ratio from two-sided Fisher’s exact test. Asterisks denote significance at Bonferroni-corrected *P* < 0.05. **b**, *C4A*-positive and *C4A*-negative genes were enriched for differentially expressed genes in SCZ brain from PsychENCODE^19^ and LIBD BrainSeq^33^. Asterisks denote significance from Fisher’s exact test at nominal *P* < 0.05. **c**, *C4A*-positive and *C4A*-negative genes were expressed in distinct cell-types. Expression-weighted cell-type enrichment (EWCE) was performed using mouse cortical/subcortical single-cell RNA-seq data^35^ and human cortical single-nucleus RNA-seq data^20^. Asterisks denote significance at FDR < 0.05. *C4A*-positive and *C4A*-negative genes are shown in red and blue, respectively.

Notably, *C4A*-positive genes were strongly enriched for co-expression modules previously shown to represent astrocyte, microglial, and NFkB signaling pathway genes. These included several canonical markers of astrocytes (e.g. *GFAP, AQP4*) and microglia (e.g. *FCGR3A, TYROBP*); critical components of the NFkB signaling pathway (e.g. *NFKB2, IL4R, RELA*); as well as known members of the classical complement pathway (e.g. *C1R, C1S*). Conversely, *C4A*-negative genes showed enrichment for neuronal and synaptic processes, stronger at higher copy number (**Supplementary Fig. 2**). These included several glutamate receptors (e.g. *GRIN2A, GRM1, GRIA3*), calcium regulators (e.g. *CAMK4, CAMTA1, CAMKK2*), and potassium channels (e.g. *KCNK1, KCNQ5, KCNIP3*). Other notable *C4A*-negative genes included the serotonin receptor *HTR2A*, the dopamine receptor *DRD1*, the major neuronal splicing regulator *NOVA1*, and the zinc transporter *SLC39A10*. These *C4A*-positive and *C4A*-negative genes were also strongly enriched for genes up- and down-regulated in SCZ brain^19,33^, respectively (**Fig. 4b**), further connecting *C4A* expression to dysregulated molecular pathways in SCZ brain.

To further refine the cell-types associated with these networks, we evaluated whether *C4A*-positive and *C4A*-negative genes were expressed in specific cell-types defined by single-cell/nucleus RNA-seq^34^. At low copy number (i.e. CN < 2), *C4A*-positive genes showed the strongest association in astrocytes, but with subsequently higher copy number, they became more broadly associated with microglia and endothelial cells (**Fig. 4c**). In contrast, *C4A*-negative genes were most highly expressed in five neuronal cell-types—cortical interneurons, pyramidal (hippocampus CA1), pyramidal (somatosensory cortex), medium spiny neurons, and striatal interneurons. Remarkably, these cell-types have all been previously shown to be enriched for SCZ GWAS signals^35^ (**Fig. 4c** and **Supplementary Fig. 3**). These findings were replicated across multiple other single-cell/nucleus RNA-seq datasets from either mouse or human brain (**Fig. 4c** and **Supplementary Fig. 4**). Taken together, these results indicate that increased *C4A* copy number is associated with brain co-expression changes leading to down-regulation of neuronal, synaptic genes—a putative transcriptomic signature of synaptic pruning.

### Sexual dimorphism of *C4A* effects in the human brain

SCZ is more prevalent in males compared with females, and recent work has identified larger effect sizes of *C4* alleles in males compared with females^36^. Although no sex differences in *C4A* expression were reported in GTEx, protein levels of *C3* and *C4* were elevated in cerebrospinal fluid (CSF) from males^36^. Here, in the independent PsychENCODE dataset, we replicated these findings, finding no sex differences in *C4A* expression in the brain (**Fig. 5a**). Notably, however, we observed a significant increase in *C4A* network size in males, consistent with larger effects in males (**Fig. 5b**; Methods). Females showed a reduction in the number of both *C4A*-positive and *C4A*-negative genes, indicating broad sex-specific effects (**Extended Data Fig. 8a**).

**Fig. 5.**
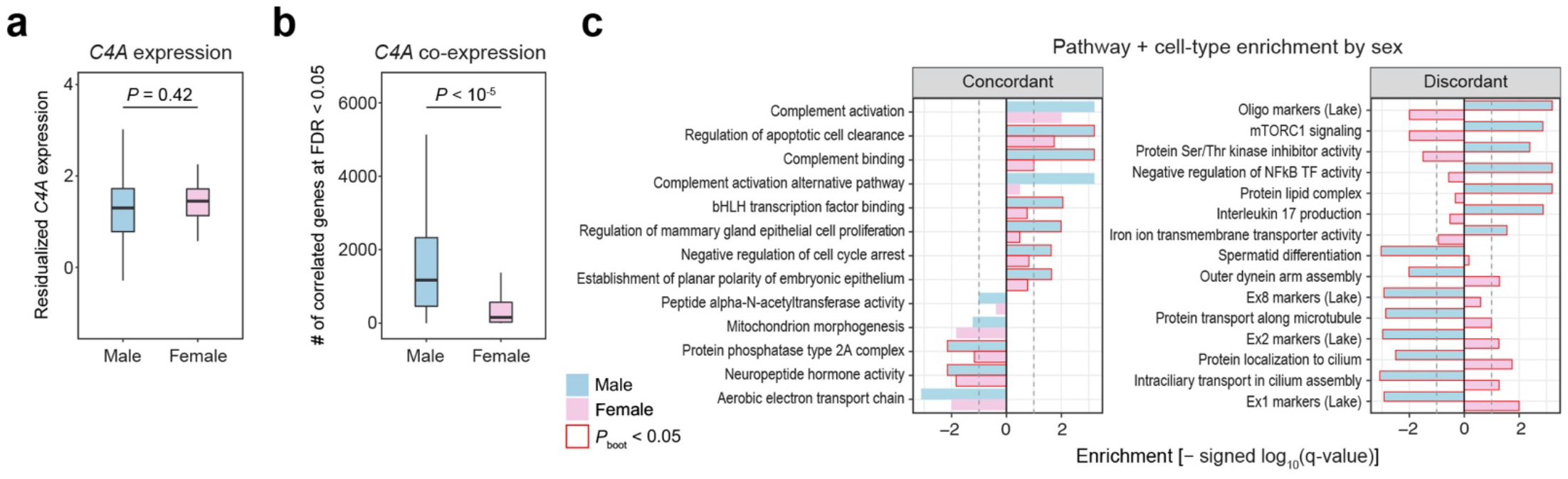
Sex differences in *C4A* co-expression highlight male-accentuated effects on mTOR signaling and neuronal cilia. **a**, Overall expression levels of *C4A* did not differ between sexes in PsychENCODE (N = 98 and 37 for male and female samples, respectively; two-sided Welch’s t-test, *P* = 0.42). **b**, Conversely, C*4A* co-expression network size was much larger in males (N = 98, 37 for males and females; permutation test, *P* < 10^−5^). Bootstrapped distributions were generated to match for sample size between sexes. **c**, To identify biological pathways and cell-types reflected by these sex-specific *C4A* co-expression patterns, we performed gene set enrichment analysis (GSEA). Genes were ranked by their *C4A* co-expression magnitude in male and female networks separately, and resulting enrichments were compared. Left, sex-concordant terms included positively associated complement activation. Right, sex-discordant terms included lipid and mTOR signaling genes as well as excitatory neuron markers and cilia-related pathways. Enrichment differences that were significant when compared to a null distribution of 10,000 random seed genes are highlighted in red. All boxplots show median and interquartile range (IQR) with whiskers denoting 1.5 × IQR.

To more systematically interrogate the neurobiological mechanisms contributing to these sexually dimorphic effects, we next sought to identify the specific pathways and cell-types that were differentially co-expressed with *C4A* across sexes. To do so, we ranked genes by the magnitude of *C4A* co-expression in males and females separately, performed gene set enrichment analysis (GSEA) on this ranked list, and compared the resulting enrichments (Methods). To ensure the robustness of these results, we further generated an empirical null distribution of enrichment differences between males and females with 10,000 randomly sampled seed genes (**Extended Data Fig. 9**; Methods). As a positive control, complement-related pathways showed concordant enrichment among *C4A*-positive genes across both sexes (**Fig. 5c**). In contrast, a number of pathways and cell-types showed significantly discordant effects across sexes. In males, *C4A*-positive genes were strongly associated with lipid and mTOR signaling genes, while these enrichments were absent in females or even showed the opposite direction of effect. Likewise, strong sex-discordant effects were observed for upper layer excitatory neuron markers^37^ (e.g. Ex1 and Ex2) and several cilia-related pathways among *C4A*-negative genes. Together, these results suggest that heightened effects of *C4A* in males may reflect distinct activation of mTOR signaling and disruption of primary cilia-related processes in excitatory neurons.

### Spatiotemporal profiles highlight frontal cortex-predominant *C4A* effects

Many biological processes occurring in the human brain are region-specific and developmentally regulated^38^. To determine whether certain regions are more susceptible to the effects of *C4A*, we next compared *C4A* network size across eight distinct brain regions from GTEx. Remarkably, we observed large regional differences with frontal and anterior cingulate cortex exhibiting the greatest degree of *C4A* co-expression (**Fig. 6a**; Methods). This result was robust to different threshold metrics (**Extended Data Fig. 8b**) and was not driven by differences in expression level across brain regions (**Fig. 6b**). These results indicate that frontal cortical regions may be particularly vulnerable to *C4A*-mediated neurobiological processes.

**Fig. 6.**
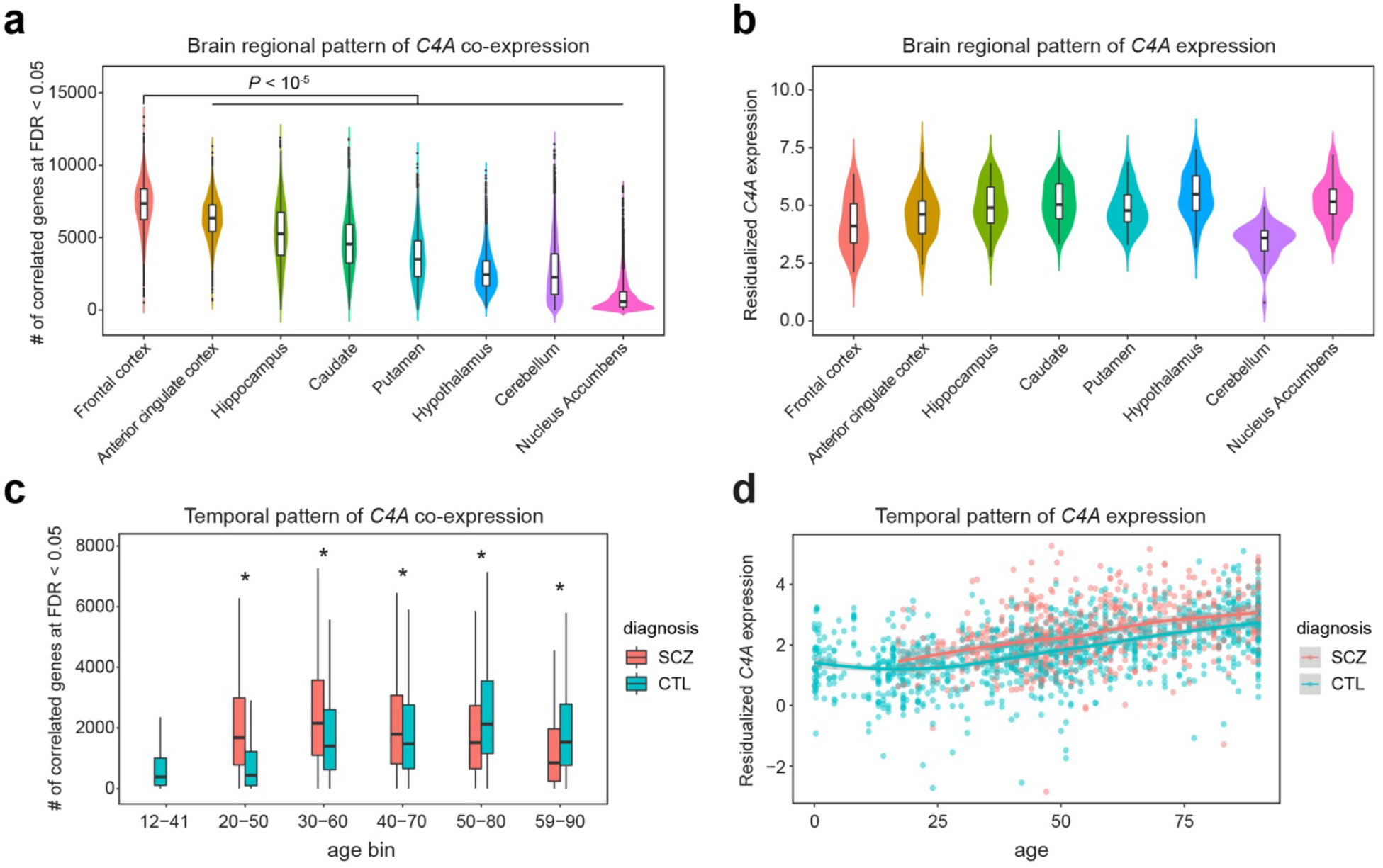
Spatiotemporal patterns of *C4A* co-expression implicate frontal cortical regions and early adult timepoints in SCZ. **a**, *C4A* exhibited the greatest degree of co-expression in frontal cortical brain areas. The plot shows the bootstrapped distribution of the number of co-expressed genes with *C4A* at FDR < 0.05 across eight different brain regions in GTEx (N = 36, 38, 45, 47, 39, 45, 39, and 45 for frontal cortex, anterior cingulate cortex, hippocampus, caudate, putamen, cerebellum, hypothalamus, and nucleus accumbens, respectively). All pairwise comparisons were statistically significant (permutation test, *P* < 10^−5^). **b**, In contrast with co-expression patterns, frontal cortical regions did not show greater *C4A* expression. The plot shows *C4A* expression in GTEx samples used for the bootstrap (N = 36, 38, 45, 47, 39, 45, 39, and 45 for frontal cortex, anterior cingulate cortex, hippocampus, caudate, putamen, cerebellum, hypothalamus, and nucleus accumbens, respectively). **c**, The temporal peak of *C4A* co-expression was earlier in SCZ cases (30- to 60-year-old window) compared to controls (50- to 80-year-old window). Bootstrapped distributions were generated across overlapping time windows using samples from PsychENCODE (N = 30, 42, 57, 68, 47, and 32 for control samples in each age bin; N = 36, 46, 55, 45, and 47 for SCZ samples). Asterisks denote significant differences in the network size between SCZ cases and controls (permutation test, *P* < 10^−5^). **d**, In contrast with co-expression patterns, *C4A* showed monotonically increasing expression across age in frontal cortex samples from PsychENCODE (N = 1730). Shown is a LOESS smooth curve with 95% confidence bands. All boxplots show median and interquartile range (IQR) with whiskers denoting 1.5 × IQR.

We next leveraged the fact that PsychENCODE contains the largest collection of uniformly processed brain samples from individuals with SCZ (N = 531) as well as neurotypical controls (N = 895) across the adult lifespan. To confer temporal resolution, we stratified samples into overlapping time windows, while controlling for *C4A* copy number, sex, and diagnosis (Methods). *C4A* co-expression reached its peak in the 50- to 80-year-old period for neurotypical controls. In comparison, a leftward age shift in co-expression peak was observed in SCZ cases (**Fig. 6c** and **Extended Data Fig. 8c**). These findings are distinct from the temporal trajectory of *C4A* expression, which increased monotonically with age (**Fig. 6d**).

### Genetic and environmental drivers of *C4A* expression alteration in SCZ brain

Finally, we sought to determine the extent to which *C4* mCNV could explain *C4A* expression alteration in SCZ brain, using frontal cortex RNA-seq data from individuals with SCZ (N = 531) and non-psychiatric controls (N = 895). As previously reported^19^, we identified strong up-regulation of *C4A* consistent with previous independent literature^9,39^. When we adjusted for *C4A* copy number, we continued to observe differential expression for *C4A* (**Fig. 7a** and **Extended Data Fig. 10**), suggesting that additional factors contribute to overexpression of *C4A* in SCZ^9,17^. Similar results were observed for several other complement system genes previously found^19^ to be differentially expressed in SCZ (**Fig. 7a** and **Extended Data Fig. 10**).

**Fig. 7.**
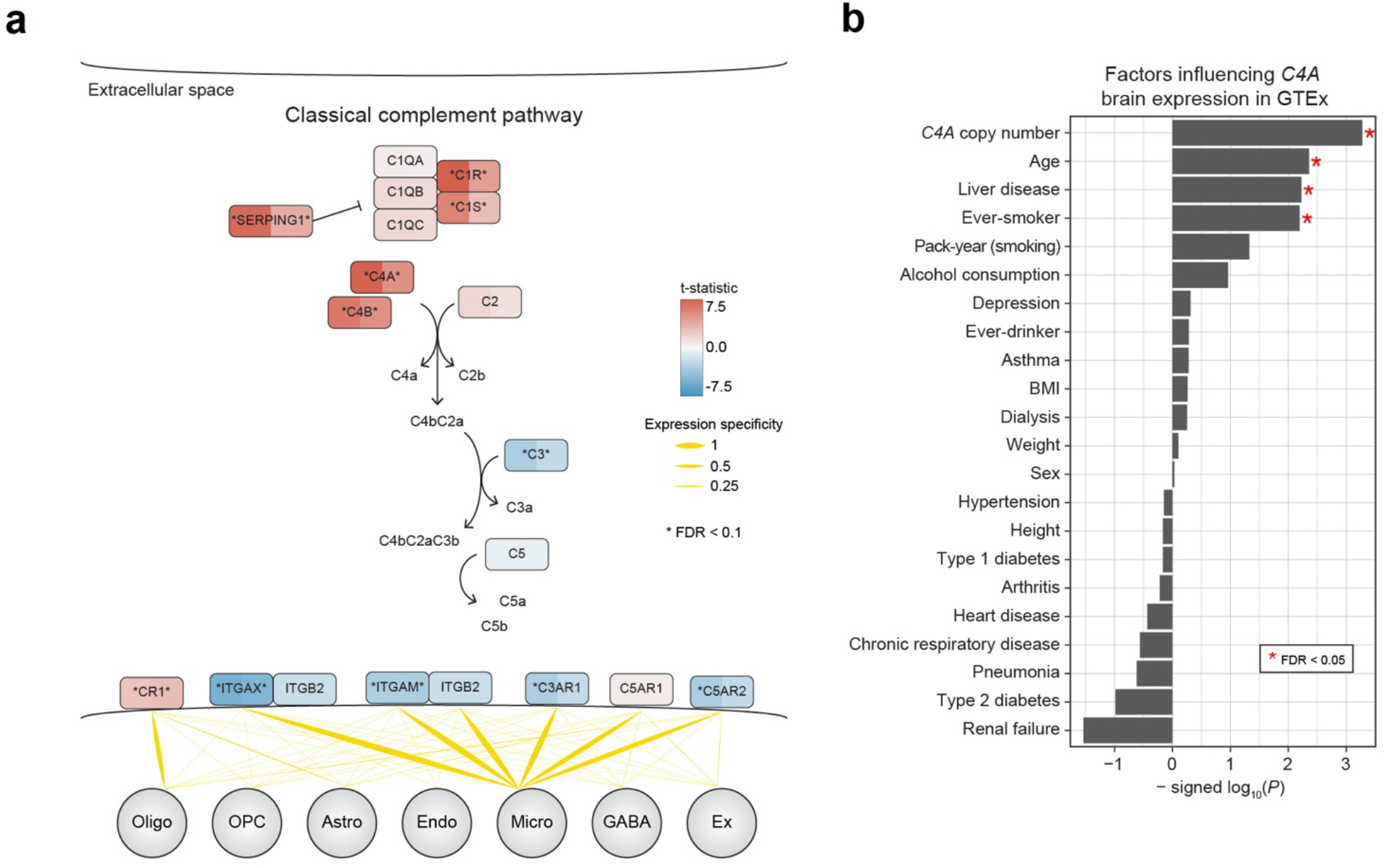
Broad, bimodal differential expression of genes within the classical complement pathway in postmortem brains from individuals with SCZ. **a**, Differential gene expression (DGE) in SCZ is shown for genes within the classical complement pathway. Early components were mostly up-regulated, whereas late components were down-regulated in SCZ. Genes are colored by DGE t-statistic on the left and t-statistic obtained while adjusting for *C4A* copy number on the right. Asterisks denote significance at FDR < 0.1. Bottom, cell-type specificity of complement receptors was calculated using snRNA-seq data from ref. ^50^. Oligo (oligodendrocyte), OPC (oligodendrocyte progenitor cell), Astro (astrocyte), Endo (endothelial), Micro (microglia), GABA (interneuron), Ex (excitatory neuron). **b**, In GTEx, we characterized the effect of documented medical comorbidities and other relevant biological covariates on brain *C4A* expression. In addition to *C4A* copy number, age, smoking, and a history of liver disease showed significant positive associations (one-sided likelihood ratio test).

To assess the specificity of these findings for SCZ, we performed an analogous analysis using frontal cortex data from individuals with bipolar disorder (BD; N = 217) and the same controls (**Extended Data Fig. 10**). Despite strong genetic and transcriptomic correlations between SCZ and BD^39^, *C4A* expression was not altered in BD, and the broader complement system exhibited minimal differential expression. This notable contrast between SCZ and BD remained when downsampling to the same number of subjects, indicating that the SCZ-BD differences were not driven by statistical power (**Extended Data Fig. 10**). Additionally, brain samples from individuals with SCZ and BD were of similar quality with respect to PMI or RIN (Welch’s t-test, *P* > 0.5) and many of the same neuroleptic medications are used to treat both conditions, indicating that these factors are unlikely to be key drivers of observed differences.

This additional component of *C4A* up-regulation in SCZ brain could be driven by other genetic factors (e.g. *trans*-eQTL) and environmental influences, or may simply represent a consequence of disease. To begin to identify potential non-genetic contributors, we turned to GTEx which has systematically compiled donor medical history. In addition to *C4A* copy number, we identified several covariates that were significantly associated with increased brain *C4A* expression—namely, age, smoking status, and a history of liver disease (**Fig. 7b**). This is notable given the substantially elevated rate of smoking in individuals with SCZ and some epidemiological evidence that smoking may increase risk for SCZ^40^. Altogether, these data support potential convergent effects of genetic (i.e. *C4* variation) and environmental (i.e. smoking) risk factors in disease risk.

## Discussion

In this study, we integrated multiple existing, large-scale genetic and transcriptomic datasets to interrogate the functional role of *C4A*—and the complement system more broadly—in the human brain and their relation to underlying core pathophysiology of SCZ. We find no evidence that the known complement system and its protein interactors are enriched for SCZ genetic signals. Using *C4A*-seeded co-expression networks, we again find that genes positively co-expressed with *C4A* show no appreciable enrichment for SCZ risk, whereas genes negatively co-expressed with *C4A* exhibit strong and specific enrichment for SCZ risk, identifying for the first time, a convergent genomic signal. These *C4A*-positive genes were associated with glial and inflammatory pathways, while *C4A*-negative genes were associated with neuronal and synaptic pathways, which is consistent with their interpretation as putative molecular correlates of synaptic pruning^9-11,16,17^. Additionally, the seeded networks expanded in size with increased genomic copy number and exhibited sexual dimorphism and spatiotemporal specificity, suggesting potential vulnerability of the adult male frontal cortex to the effects of *C4A*. Overall, these results highlight convergence of SCZ polygenic effects and indicate that synaptic processes—rather than the complement system—are the driving force conferring SCZ risk (**Fig. 8**).

**Fig. 8.**
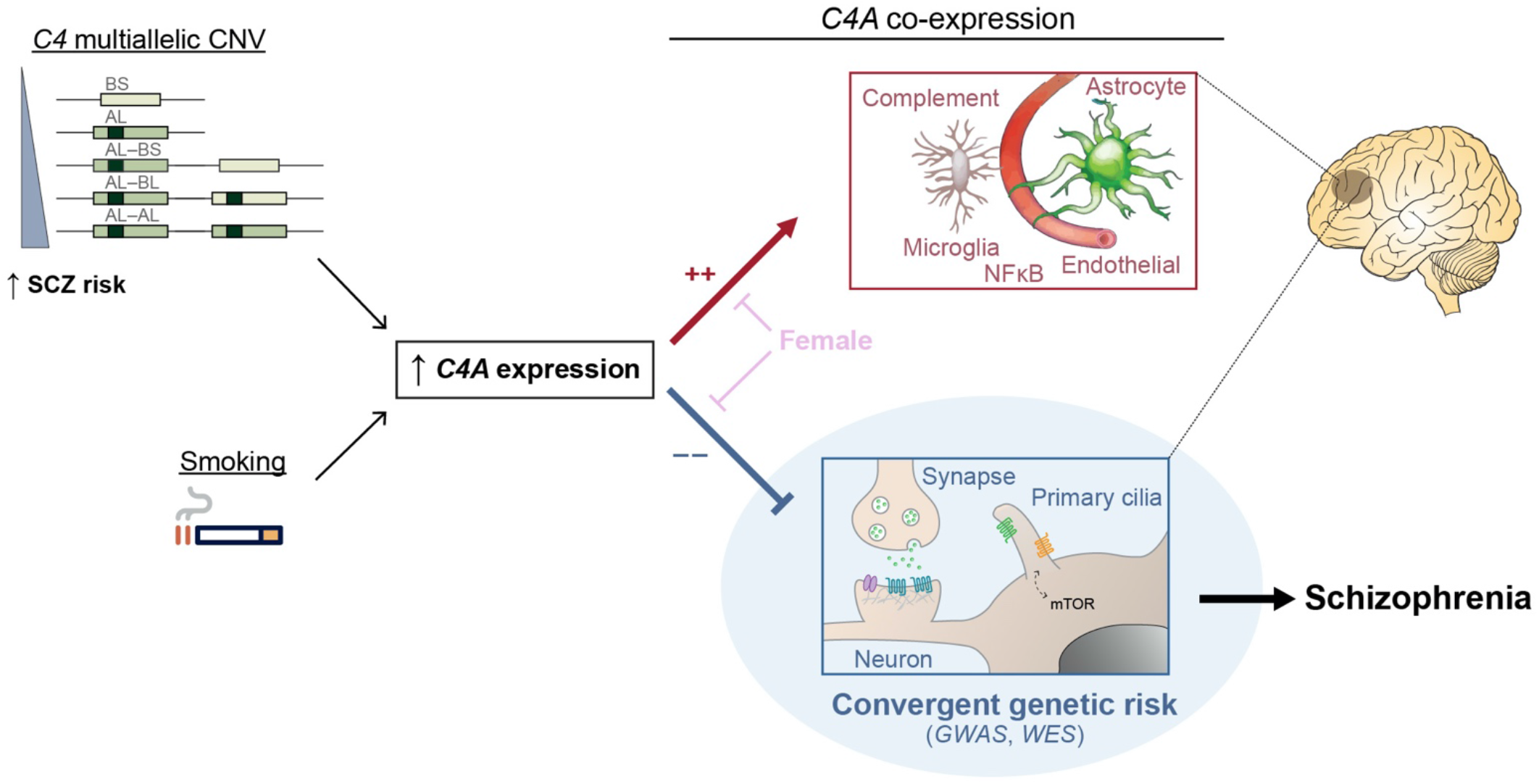
A model of the functional role of *C4A* in SCZ pathogenesis. mCNV of *C4* genes as well non-genetic factors such as smoking influence *C4A* expression. *C4A* expression is positively associated with glial and inflammatory processes and negatively associated with neuronal and synaptic processes, which in turn are enriched for SCZ genetic signals.

We first observed that SCZ genetic risk is not enriched among complement system genes—despite testing multiple classes of genetic variation (i.e. GWAS, rare variants, large recurrent CNVs), using multiple statistical methods with varying genomic window sizes, and expanding the annotation to include high-confidence PPIs or *C4A*-positive genes. This was surprising given the integral role of *C4A* in the complement system^10^, the strength of the *C4A* association^9^, and the high level of polygenicity observed in SCZ^4,5^. This does, however, comport with recent East Asian SCZ GWAS results^41^, which did not observe an MHC association, despite a genetic correlation of 0.98 with European GWAS results. These findings imply that dysregulation of the complement system is neither necessary nor sufficient for the development of SCZ and fit with an alternative explanation that *C4A* may be more associated with the progression or severity of illness. Moreover, the logical extension of these observations predicts that drugs targeting this pathway are unlikely to be a panacea.

How then does *C4A* impart risk for SCZ? We reasoned that the functional role of *C4A* in the human brain may not be well captured by manually curated gene sets and pathway annotations, which are often incomplete. To address this, we leveraged co-expression networks and subsequent guilt-by-association to generate an unbiased, human brain-relevant functional annotation for *C4A*. As expected, *C4A*-positive genes capture the known complement system and reflect inflammatory processes, including astrocyte, microglial, and NFkB signaling pathways—all of which are dysregulated in SCZ brain^19^, but none of which show an appreciable enrichment for genetic risk. Similar changes have been observed in other neuropsychiatric disorders^19,39^ and may reflect environmental influences (e.g. smoking) or represent the consequence of a more proximal (e.g. synaptic) pathology. In contrast, *C4A*-negative genes reflect dysregulated neuronal and synaptic pathways, exhibiting strong genetic enrichment for SCZ. Notably, the network size and connectivity expand substantially with increased *C4A* copy number, indicating that *C4A* plays more of a driver role with increasing genetic risk for SCZ.

We find that *C4A* CNV is strongly associated with—but does not fully explain—the observed *C4A* up-regulation in SCZ brain. Similarly, although our results suggest that *C4A*-mediated SCZ risk occurs through synaptic mechanisms rather than complement signaling, several additional complement system genes exhibit differential expression in SCZ, even when controlling for *C4A* copy number. This included up-regulation of early components (e.g. *C1R, C1S*), but also significant down-regulation of downstream components including known complement receptors (e.g. *ITGAM, ITGAX, C3AR1, C5AR2*). We hypothesize that some of these observed transcriptomic alterations reflect a compensatory response to synaptic dysfunction, as *C4A* up-regulation has also been observed in ASD brain^19^, despite not being considered a genetic risk factor. Additionally, we find that brain *C4A* expression is elevated with smoking. Intriguingly, smoking is associated with diffuse, dose-dependent cortical thinning^42^ and there is epidemiological evidence supporting a directional effect of smoking on SCZ risk^40^, although confounding factors (e.g. cannabis use) likely also contribute^43^. Overall, these results highlight a neurobiological mechanism through which genetic and environmental risk factors converge and contribute to SCZ risk.

Finally, comparison of the network size provided additional insights into the spatiotemporal and sex-specific effects of *C4A*. Males showed greater degree of *C4A* co-expression, despite comparable *C4A* expression level across sexes, which is consistent with larger effects of *C4A* alleles in males relative to females^36^. Compared to its female counterpart, male *C4A*-seeded network showed greater activation of lipid and mTOR signaling pathways as well as greater disruption of cilia-related processes and excitatory neuron markers (**Fig. 8**). Both mTOR signaling and primary cilia are known to be critical regulators of neurogenesis and synaptic pruning^44-47^. Primary cilia, the solitary microtubule-based structure present in most neurons, glia, and their progenitors, also serve as a major hub for signaling pathways, including mTOR, Sonic hedgehog (Shh), Wnt, autophagy, and ubiquitin-proteasome system^48^, several of which have intriguing links to SCZ and other neurodevelopmental disorders^29,49^ that warrant further experimental investigation. Together, these findings highlight several potential mechanisms underlying greater disease vulnerability in males. Importantly, these observations are only evident through analysis of co-expression, rather than expression patterns alone, demonstrating the importance of this approach.

We note that several important questions remain for *C4A* in relation to SCZ. Although we identify *C4A*-specific interaction with *C4A* copy number variation, *C4A* and *C4B* co-expression partners are highly similar in general, making it difficult to disambiguate the effects of *C4A* from *C4B*. Further work in characterizing the biochemical properties of C4 proteins in the human brain is necessary to fully elucidate the mechanism through which *C4A* exerts larger effects in SCZ. In addition, human cell-types that express *C4A* in either physiology or pathophysiology remain unclear, due to dropout events in single-cell/nucleus RNA-seq. Although *C4A*-positive genes at low copy number (i.e. CN < 2) show strong and selective enrichment for astrocytes, and expression specificity of *C4A* is similarly the highest in astrocytes according to various mouse single-cell RNA-seq datasets^35^, this remains to be validated for humans in future studies. Spatiotemporal resolution is also relatively restricted in this study, since the scope of our analyses is inherently limited to the range of available functional genomic resources, and our use of post-mortem samples is limited to retrospective analyses which cannot directly infer causal relationships. As larger and more diverse samples spanning all SCZ-relevant regions (e.g. striatum) and developmental time points become available, spatiotemporal specificity will undoubtedly improve. Likewise, as human brain genomic panels increase in size, we anticipate additional insights to be gained from distal genetic regulators (e.g. *trans*-eQTL) of *C4A*. Lastly, model systems capable of fully recapitulating postnatal neuronal-glial interactions in the human frontal cortex will be necessary for experimental validation.

## Acknowledgments

This work was supported by the Simons Foundation Bridge to Independence Award (MJG), the National Institute of Mental Health (R01MH121521 to M.J.G.; R01MH123922 to M.J.G.; P50HD103557 to M.J.G.; K00MH119663 to L.M.H.; T32MH073526 to M.K.), and the UCLA Medical Scientist Training Program (T32GM008042 to M.K.). The authors thank Gil Hoftman and members of the Gandal Lab for critical comments. Data were generated as part of the PsychENCODE Consortium, supported by: U01MH103392, U01MH103365, U01MH103346, U01MH103340, U01MH103339, R21MH109956, R21MH105881, R21MH105853, R21MH103877, R21MH102791, R01MH111721, R01MH110928, R01MH110927, R01MH110926, R01MH110921, R01MH110920, R01MH110905, R01MH109715, R01MH109677, R01MH105898, R01MH105898, R01MH094714, P50MH106934, U01MH116488, U01MH116487, U01MH116492, U01MH116489, U01MH116438, U01MH116441, U01MH116442, R01MH114911, R01MH114899, R01MH114901, R01MH117293, R01MH117291, R01MH117292 awarded to: Schahram Akbarian (Icahn School of Medicine at Mount Sinai), Gregory Crawford (Duke University), Stella Dracheva (Icahn School of Medicine at Mount Sinai), Peggy Farnham (University of Southern California), Mark Gerstein (Yale University), Daniel Geschwind (University of California, Los Angeles), Fernando Goes (Johns Hopkins University), Thomas Hyde (Lieber Institute for Brain Development), Andrew Jaffe (Lieber Institute for Brain Development), James Knowles (University of Southern California), Chunyu Liu (SUNY Upstate Medical University), Dalila Pinto (Icahn School of Medicine at Mount Sinai), Panos Roussos (Icahn School of Medicine at Mount Sinai), Stephan Sanders (University of California, San Francisco), Nenad Sestan (Yale University), Pamela Sklar (Icahn School of Medicine at Mount Sinai), Matthew State (University of California, San Francisco), Patrick Sullivan (University of North Carolina), Flora Vaccarino (Yale University), Daniel Weinberger (Lieber Institute for Brain Development), Sherman Weissman (Yale University), Kevin White (University of Chicago), Jeremy Willsey (University of California, San Francisco), and Peter Zandi (Johns Hopkins University). The Genotype-Tissue Expression (GTEx) Project was supported by the Common Fund of the Office of the Director of the National Institutes of Health, and by NCI, NHGRI, NHLBI, NIDA, NIMH, and NINDS.

## Author contributions

M.K. and M.J.G. planned the study and wrote the paper. M.K. performed primary analyses with J.R.H., P.Z., L.M.H., L.M.O.L., L.dlT-U. and M.J.G. providing additional input. L.W. and L.P.C. validated the *C4* imputation pipeline. All authors read and approved the final manuscript.

## Methods

### Annotation of the complement system and its protein-protein interactions (PPIs)

We compiled a list of 57 genes annotated as part of the complement system in the HUGO Gene Nomenclature Committee (HGNC) database (genenames.org)^51^. Of these, 42 genes were found to be expressed in the PsychENCODE RNA-seq data, after filtering for genes with TPM > 0.1 in at least 25% of samples^19^. Those missing (n = 15 genes) due to low expression included: *C6, C8A, C8B, C9, FCN2, MBL2, C4BPA, C4BPB, CFHR1, CFHR2, CFHR3, CFHR4, CFHR5, F2*, and *CR2*. The annotation was also expanded by including high-confidence human PPIs for the complement system with score > 0.7 from the InWeb3 database^27^ (n = 57 + 488 = 545 genes) (**Supplementary Table 1**).

### Evaluation of the complement system for common variant association

We evaluated the proximity of the complement components to genome-wide significant loci from two recent SCZ GWAS studies^4,5^. Four genes (*C4A, C4B, CFB, C2*) were within the MHC region. Excluding the MHC, nine genes (*SERPING1, CLU, CSMD1, CD46, CD55, CR1, CR2, C4BPA*, and *F2*) were within 1 Mb of GWAS loci. These genes were subsequently assessed for Hi-C interactions in fetal and adult brain^23^ and significance from SMR method using brain and whole blood eQTL panels from PsychENCODE^19^ and eQTLGen^52^, respectively.

### Stratified LD score regression (sLDSC)

sLDSC^25^ was used to test whether a gene set of interest is enriched for SNP-based heritability in various phenotypes (i.e. disease and trait)^5,53-70^. SNPs were assigned to custom gene categories if they fell within ±100 kb of a gene in the set. For the complement system, we also tested a range of window sizes (±1 kb to 1 Mb) around each gene. These categories were then added to a full baseline model that includes 53 functional categories capturing a broad set of genomic annotations. The MHC region was excluded from all analyses by default. Enrichment was calculated as the proportion of SNP-based heritability accounted for by each category divided by the proportion of total SNPs within the category. Significance was assessed using a block jackknife procedure, followed by Bonferroni correction for the number of phenotypes tested.

### MAGMA

MAGMA (v1.07b)^26^ was used to assess enrichment of SCZ GWAS signals among the complement system. An annotation step was first performed in which SNPs in a specified window surrounding each gene were combined, while accounting for linkage disequilibrium (LD). We tested several window sizes ranging from ±0 kb to 100 kb, and LD was calculated using the European panel of 1000 Genomes Project^71^. A competitive gene-level analysis was then performed using the complement annotations defined above.

### Rare variant enrichment

Multiple gene sets were assessed for enrichment of rare variants identified in neurodevelopmental disorders. These included: ∼100 high-confidence autism spectrum disorder (ASD) risk genes harboring rare *de novo* variants^72,73^; ASD risk genes harboring rare inherited variants^74^; genes harboring recurrent *de novo* copy number variants associated with ASD or SCZ, as compiled in ref. ^39^; genes harboring an excess of rare exonic variants in ASD, SCZ, intellectual disability (ID), developmental delay (DD), and epilepsy as assessed through an extended version of transmission and *de novo* association test (extTADA)^75^; syndromic and highly ranked (1 and 2) genes from SFARI Gene database (https://gene.sfari.org/); genes harboring disruptive and damaging ultra-rare variants (dURVs) in SCZ cases^30^; a list of high-confidence epilepsy risk genes compiled in ref. ^76^; risk genes for developmental disorders harboring rare *de novo* variants^77^; and ten high-confidence SCZ risk genes harboring rare exonic variants as identified by the SCHEMA consortium^29^. For binary gene sets, statistical enrichment analyses were performed using logistic regression, correcting for linear- and log-transformed gene and transcript lengths as well as GC content. For dURVs, a two-step procedure was used, first creating a logistic regression model for genes harboring dURVs in controls and a second model for those affected in cases and controls. The likelihood ratio test (LRT) was used to assess significance. For SCHEMA and extTADA gene sets, the -log_10_-transformed *P* value and posterior-probability (PP) was used, respectively, in place of binary annotation in the above logistic regression model. All results were FDR-corrected for multiple comparisons.

### The PsychENCODE brain genomic dataset

Genotype array and frontal cortex RNA-seq data from Freeze 1 and 2 of PsychENCODE were obtained from www.doi.org/10.7303/syn12080241. This consisted of uniformly processed data from six studies: BipSeq, LIBD_szControl, CMC_HBCC, CommonMind, BrainGVEX, and UCLA-ASD (see Table S1 and Fig. S33 in ref. ^20^). Genotype data for these individual studies were previously harmonized^20^ through phasing and imputation with the Haplotype Reference Consortium (HRC) reference panel. We used post-QC RNA-seq data that were fully processed, filtered, normalized, and extensively corrected for all known biological and technical covariates except the diagnosis status (see Materials/Methods and Fig. S3 in ref. ^19^). Of note, RNA-seq reads were previously aligned to the hg19 reference genome with STAR 2.4.2a and gene-level quantifications calculated using RSEM v1.2.29. Genes were filtered to include those with TPM > 0.1 in at least 25% of samples^19^. The same expression data were used for all downstream analyses unless otherwise stated.

### Imputation of *C4* structural alleles

The *C4* locus harbors multiallelic CNV (mCNV), where human *C4* encoded by two genes (*C4A* and *C4B*) can exist in different combinations of copy numbers. The two paralogs are defined based on four amino acid residues in exon 26, which are thought to alter binding affinities for distinct molecular targets. Either paralog can also contain a human endogenous retroviral insertion (*C4*-HERV) in intron 9, which then functions as an enhancer and preferentially increases *C4A* expression^9^. Recent work demonstrated that four common *C4* structural alleles are in linkage disequilibrium (LD) with nearby SNPs^9^ and hence can be accurately imputed from genotype array data. Accordingly, we imputed *C4* alleles in six studies from PsychENCODE separately using Beagle4.1^78^ with a custom HapMap3 CEU reference panel as described^9^. We began with the HRC imputed genotype data and filtered for high-quality SNPs by setting the R2 > 0.3 threshold from Minimac3. We restricted imputation and subsequent downstream analyses to samples of European ancestry (N = 812) based on genetic principal component analysis with the 1000 Genomes Project reference panel^71^ (**Extended Data Fig. 1**). There was an overlap of individuals in BipSeq, LIBD_szControl, and CMC_HBCC studies, which used different SNP genotyping platforms (see Table S1 in ref. ^20^). For these duplicate samples, the concordance rate of imputation result was high (N = 181/204 individuals with matching result), indicating robust *C4* imputation. For 23 samples with discordant imputation results, we calculated the average dosage for each structural allele and inferred the most likely pair of structural alleles.

### Effect of *C4* variation on gene expression

Inferred copy number of *C4* structural elements (*C4A, C4B*, and *C4*-HERV) based on the imputed *C4* alleles was associated with *C4A* and *C4B* RNA expression using a linear model. Both best-guess copy number and probabilistic dosage were tested for association, which yielded an analogous result. As shown previously^9,19,79^, *C4A* expression was strongly associated with *C4A* copy number (R = 0.37, *P* = 2.8 × 10^−27^) and *C4*-HERV copy number (R = 0.33, *P* = 7.9 × 10^−22^), but not with *C4B* copy number (R = -0.03, *P* = 0.39; **Extended Data Fig. 5**). Likewise for *C4B*, expression was associated with corresponding gene dosage (R = 0.12, *P* = 3.8 × 10^−4^), but not with *C4A* copy number (R = -0.05, *P* = 0.15) or *C4*-HERV copy number (R = -0.05, *P* = 0.17).

### Construction of *C4A*-seeded networks

To ensure imputation quality and thereby draw robust biological inference, we restricted our network analyses to samples with average imputed probabilistic dosage > 0.7 (N = 552/812). Most studies had high probabilistic dosage, except BrainGVEX and UCLA-ASD. In the case of BrainGVEX, this was because there were many missing SNPs in the vicinity of *C4* locus. This filtering step hence removed most samples with low-quality imputation from BrainGVEX and UCLA-ASD. Neurotypical control samples with diploid *C4A* CN = 2 (N = 145/552) (**Extended Data Fig. 2**) were then used to generate a *C4A*-seeded network by calculating pairwise PCC between *C4A* and 25,774 features, which included 16,541 protein-coding and 9,233 noncoding genes based on Gencode v19 annotations (**Supplementary Table 2**). To test whether this network is enriched for the known complement components than can be expected by chance, we randomly sampled 10,000 seed genes and generated 10,000 seeded networks. For each network, genes positively correlated with the seed gene at FDR < 0.05 were assessed for overlap with the annotated complement system (n = 57 genes), while genes negatively correlated with the seed gene at FDR < 0.05 were assessed for overlap with genes annotated within the SynGo database^31^ (n = 1,103 genes; **Extended Data Fig. 4**).

To capture broad genetic effects of *C4A* CNV on *C4A* co-expression, we stratified PsychENCODE samples into three CNV groups (i.e. CN < 2, CN = 2, and CN > 2). For control samples, there were at least 54 samples in each group (**Extended Data Fig. 2**). To account for uneven sample sizes, we used 10,000 bootstrapping replicates to downsample to 50 samples across each group. We calculated PCC in every iteration as above and eventually took the median PCC and its corresponding *P* value. Generated using only the control samples, these networks were not influenced by case-control status and disease-associated confounding factors (e.g. medication and RNA degradation effects). Additionally, the control samples were balanced in covariates such as age, RIN, postmortem interval (PMI), brain pH, and sex (**Supplementary Fig. 5**).

To maximize sample size and hence power to detect significant co-expression, particularly for rarer *C4A* CNV groups (i.e. CN < 2 and CN > 2), we also constructed the seeded networks by using every sample that passed the above quality control (N = 552). Combining all samples irrespective of the diagnosis status led to a minimum of 109 samples in each CNV group (**Extended Data Fig. 2**), allowing us to generate the networks with bootstrap by downsampling to 100 samples. Such all-sample networks yielded analogous results to control-only networks in terms of the network expansion with respect to *C4A* CNV, effect sizes of *C4A* co-expression, and the patterns of pathway, cell-type, and genetic enrichments (**Supplementary Fig. 6**). Given the robustness of these network findings, we present results from all-sample networks. For visualization of the *C4A*-seeded networks, a hard-threshold of PCC > 0.5 and FDR < 0.05 was applied. All network plots were drawn using *igraph* and *ggplot2* packages in R.

### The GTEx brain genomic dataset

GTEx v7 was used for external replication^21^. We downloaded the GTEx genotype data from dbGaP (accession phs000424.v7.p2) and imputed *C4* alleles in samples of European ancestry according to genetic principal component analysis. We obtained transcript-level counts from www.gtexportal.org and derived gene-level counts using *tximport* package in R. Briefly, RNA-seq reads were aligned to the hg19 reference genome with STAR 2.4.2a and transcript-level counts quantified with RSEM v1.2.22. We started with samples and features that were used for GTEx eQTL analyses. We then dropped samples from non-brain tissues and tissues with different sample preparation (i.e. cortex and cerebellar hemisphere). We also dropped samples with a history of disease possibly affecting the brain prior to filtering for features with CPM > 0.1 in at least 25% of samples. Gene-level counts were then normalized using TMM normalization in edgeR and log_2_-transformed to match PsychENCODE. Each brain region was then assessed for outlier samples, defined as those with standardized sample network connectivity Z scores < -3, which were removed. These quality control steps resulted in 20,765 features based on Gencode v19 annotations and 920 samples across ten brain regions, out of which 540 samples were imputed for *C4* alleles.

We next regressed out biological and technical covariates except region and subject terms using a linear mixed model via *lme4* package in R. We entered region, age, sex, 13 seqPCs (top 13 principal components of sequencing QC metrics from RNA-SeQC), RIN, ischemic time, interval of onset to death for immediate cause, Hardy Scale, body refrigeration status as fixed effects and subject as a random intercept term. To evaluate the relationship between several non-genetic factors and *C4A* gene expression, we added 3 genetic PCs, brain pH, and a covariate of interest (e.g. BMI, weight, height, smoking status, or drinking status) as fixed effects to the above model. Significance was assessed by the likelihood ratio test (LRT) of the full model with the effect in question against the null model without the effect in question.

Due to the relatively limited sample size of GTEx (i.e. less than 10 samples for CN < 2 and CN > 2 in each brain region), we focused on samples with two *C4A* copy number in subsequent analyses. We constructed a *C4A*-seeded network using frontal cortical samples (N = 36) and combined this with the above PsychENCODE control-only network (N = 145) using the Olkin-Pratt (OP) fixed-effect meta-analytical approach as implemented in *metacor* R package.

### Interaction of *C4A* copy number with *C4A* expression

The specificity of the *C4A*-seeded network expansion with respect to *C4A* CNV was evaluated statistically via multiple linear regression. We tested for an interaction term between *C4A* copy number variation and *C4A* gene expression on other gene targets transcriptome-wide (i.e. 25,774 brain-expressed genes). Given that *C4A* copy number and *C4B* copy number are negatively correlated with one another (Pearson’s R = -0.41, *P* = 1.3 × 10^−23^), both terms were included in our regression. The model we tested was: gene_*j*_ ∼ (*C4A* CN + *C4B* CN) × *C4A* expr, where the subscript *j* refers to the expression of gene *j* (**Extended Data Fig. 7**). To determine how these results compare to what would be expected by chance, we replaced *C4A* expression in the above model by a randomly selected gene and calculated the number of times the interaction term was significant. We repeated this until we had randomly sampled 10,000 genes, and the empirical *P* values for *C4A* and *C4B* expression were subsequently calculated (*P* = 10^−4^ and 0.11, respectively).

### Pathway enrichment

For pathway enrichment, we focused on genes co-expressed with *C4A* at FDR < 0.05. Enrichment for GO terms was performed using *gProfileR* v0.6.7 package in R with strong hierarchical filtering (**Supplementary Fig. 1**). Only pathways containing less than 1,000 genes and more than 10 genes were assessed. Background was restricted to brain-expressed genes and an ordered query was used, ranking genes by correlation with *C4A*. Overlap with PsychENCODE WGCNA modules^19^ was assessed using Fisher’s exact test, followed by Bonferroni correction for multiple testing (**Supplementary Fig. 2**). The same gene sets were finally assessed for overlap with differentially expressed genes (DEG) in SCZ brain from PsychENCODE^19^ and LIBD BrainSeq Phase II^33^. For PsychENCODE, DEG at FDR < 0.05 were tested, while for LIBD BrainSeq, DEG at FDR < 0.1 were tested.

### Expression-weighted cell-type enrichment (EWCE)

We used 10,000 bootstrapping replicates for EWCE with genes co-expressed with *C4A* at various FDR thresholds (**Supplementary Figs. 3-4**). Briefly, EWCE statistically evaluates whether a gene set of interest is expressed highly in a given cell-type than can be expected by chance. Z-score is estimated by the distance of the mean expression of the target gene set from the mean expression of bootstrapping replicates^34^. We downloaded pre-computed expression specificity values for several single-cell/nucleus RNA-seq data from http://www.hjerling-leffler-lab.org/data/scz_singlecell/. For independent single-nucleus RNA-seq datasets from refs. ^20,50^, we processed and computed the expression specificity metric of each gene as described^34,35^.

### Sex differences in *C4A* co-expression

As there were fewer female than male samples in PsychENCODE, we combined the control samples with two *C4A* copy number in the 12-to 80-year-old period for each sex separately. The resulting samples were balanced in age (Welch’s t-test, *P* = 0.70; Wilcoxon rank-sum test, *P* = 0.54). We then tested for sex differences in *C4A* co-expression using bootstrapping to match the sample size (37 samples + 10,000 iterations). To identify pathways and cell-types differentially co-expressed with *C4A* across sex, we ranked genes by the magnitude of *C4A* co-expression in male and female samples separately. This ranked list was then used for gene set enrichment analysis (GSEA)^80^ using the *clusterProfiler* R package. The union of GO and Hallmark gene sets from the MSigDB collections (C5 + H v7.1)^81^, gene sets from SynGO^31^, and the human brain cell-type markers defined in ref. ^37^ were tested for enrichment. To assess significance of GSEA results, we randomly sampled 10,000 seed genes. For each seed gene, we calculated male and female-specific co-expression and performed GSEA as above. The difference in normalized enrichment score (NES) between sexes was used as the test statistic. The empirical *P* value for each gene set was subsequently calculated by comparing the rank of this difference for *C4A* to the empirical null distribution of the test statistic from randomly sampled seed genes (**Extended Data Fig. 9**).

### Spatial resolution of *C4A* co-expression

To ensure the robustness of co-expression results, we focused on eight brain tissues from GTEx that had at least 35 samples with two *C4A* copy number^82,83^. As the number of samples varied across brain regions (i.e. N = 36, 38, 45, 47, 39, 45, 39, and 45 for frontal cortex, anterior cingulate cortex, hippocampus, caudate, putamen, cerebellum, hypothalamus, and nucleus accumbens, respectively), we used 10,000 bootstrapping replicates to downsample to 36 samples. In each iteration, we calculated PCC between *C4A* and every other gene and estimated the number of significantly co-expressed genes at FDR < 0.05. Other threshold metrics were tested as well, which gave similar results (**Extended Data Fig. 8**). We did not control for other biological covariates such as age and sex to maximize sample size and also because they were not significantly different across brain regions (one-way ANOVA, *P* = 0.99; Fisher’s exact test, *P* = 0.95).

### Temporal resolution of *C4A* co-expression

As our analyses suggest that *C4A* copy number variation exhibits strong genetic effects on *C4A* co-expression, we controlled for *C4A* copy number by focusing on samples with two *C4A* copy number in PsychENCODE. In order to reduce other sources of bias such as sex and diagnosis, we only used male samples and performed separate analyses for controls and SCZ cases. We divided the samples by six overlapping time windows and calculated the number of co-expressed genes for *C4A* in each time period with bootstrap (30 samples + 10,000 iterations). Here, we note relatively limited sample size and crude time windows post-stratification of the PsychENCODE dataset in order to control for potential confounding factors.

### Differential expression of the complement system

Differential gene expression of the complement was calculated using a linear mixed model via *nlme* package in R as previously reported^19^. We repeated this analysis by randomly downsampling SCZ samples to match the sample size of BD. We additionally performed several conditional analyses by adjusting for *C4A* expression and/or *C4A* copy number (**Extended Data Fig. 10**). As *C4* alleles were imputed in only the samples of European ancestry, a subset of PsychENCODE was used for such conditional analyses.

### Statistics and reproducibility

No statistical methods were used to pre-determine sample sizes, but our study makes use of the largest publicly available genomic dataset of postmortem human brains^19,20^. Even after stratifying samples by imputed *C4A* copy number, this sample size was sufficient^82,83^ to detect significant gene co-expression, as we observed. Randomization and blinding were not possible due to the study being retrospective and observational. Accordingly, subject-level covariates were used to account for variation in gene expression as well as to remove unwanted confounding effects. We downloaded and uniformly processed the independent data from the GTEx project for external replication of PsychENCODE findings. Overall co-expression pattern and subsequent cell-type, pathway, and genetic enrichment results were replicated. We did not attempt to replicate the network expansion findings due to the small sample size of GTEx for rare copy number variant groups. For differential expression analyses across sex and case-control status, normalized gene expression was assumed to follow normal distribution, but this was not formally tested. Effects of genetic and environmental factors on gene expression were also assessed using a linear model. Additional details for statistical analyses are provided in relevant sub-sections of the Methods.

## Data availability

PsychENCODE raw genotype and RNA-seq data that support the findings of this study are available at www.doi.org/10.7303/syn12080241. Processed PsychENCODE summary-level data are available at Resource.PsychENCODE.org. GTEx genotype and RNA-seq data used for the analyses described in this manuscript were obtained from: the GTEx Portal (www.gtexportal.org) and dbGaP accession number phs000424.v7.p2.

## Code availability

The code used to perform bioinformatic analyses are available at: https://github.com/gandallab/C4A-network.

