## Supplementary Figures for "Brain gene co-expression networks link complement signaling with convergent synaptic pathology in schizophrenia"

### Supplementary Information

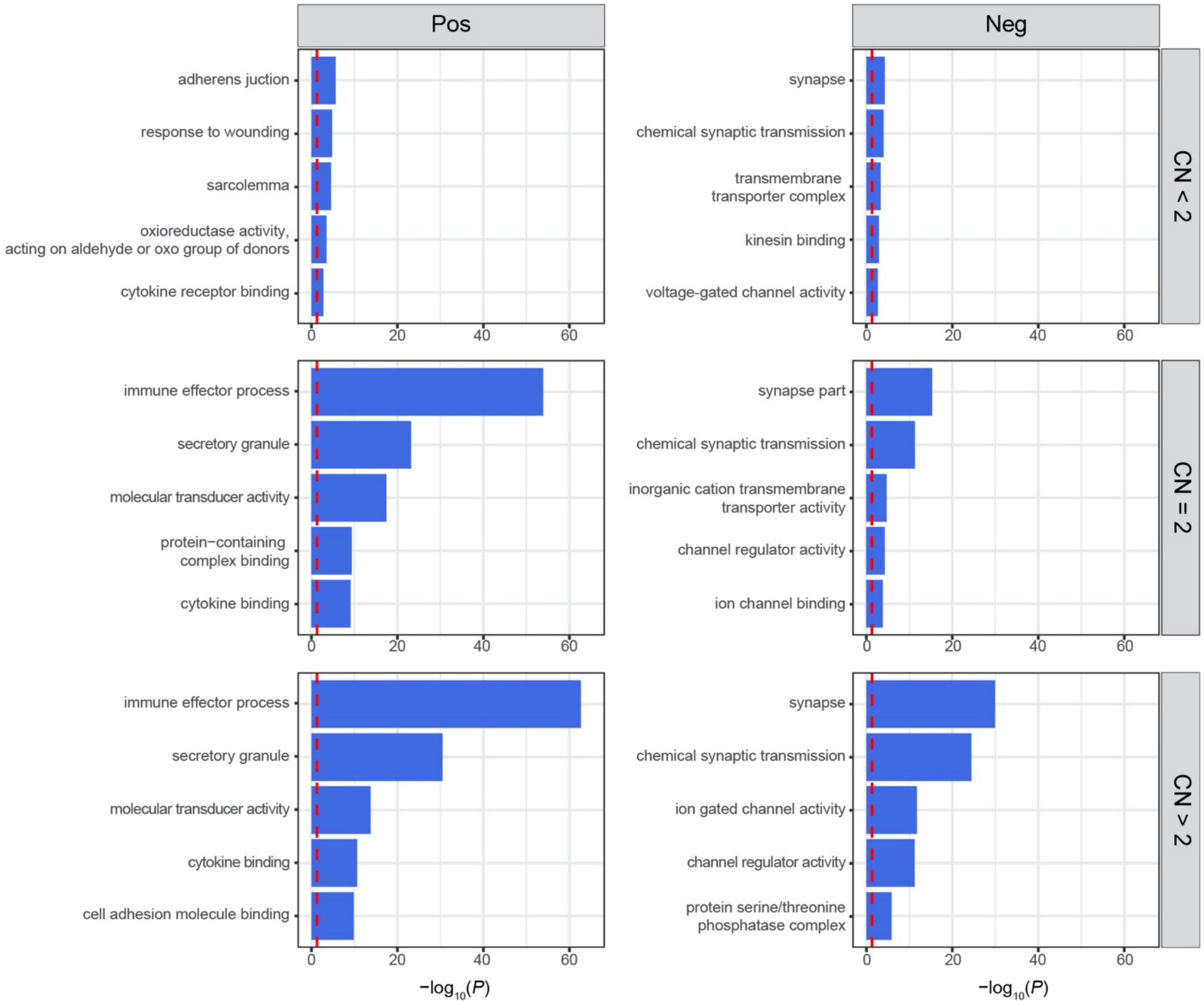


**Supplementary Figure 1. Enrichment for distinct GO terms among *C4A*-positive and *C4A*-negative genes.** Gene sets obtained from the seeded networks at FDR < 0.05 were used for pathway enrichment analyses. Top five GO terms with the highest enrichment are shown. The red dotted line denotes FDR-adjusted *P* value at 0.05.

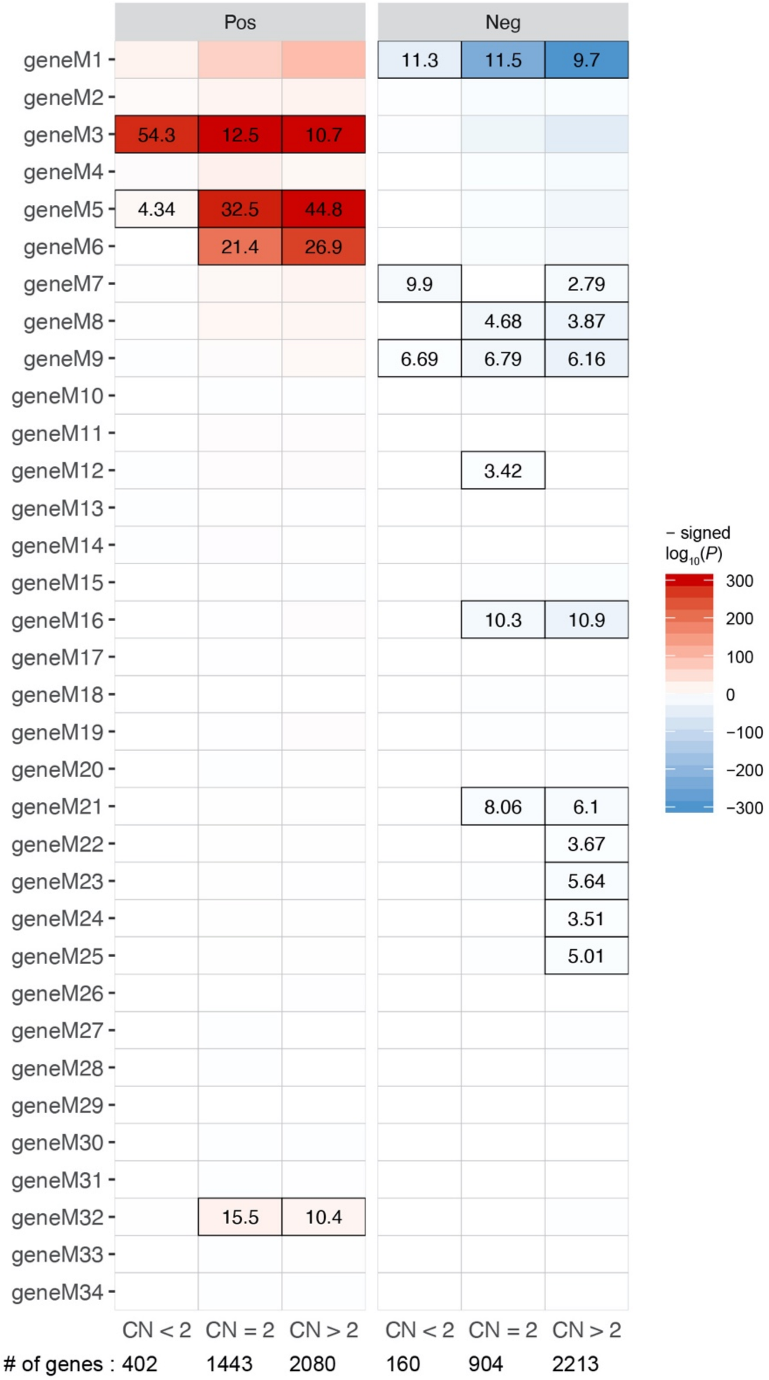


**Supplementary Figure 2. Enrichment for distinct WGCNA modules among *C4A*-positive and *C4A*-negative genes.** Gene sets obtained from the seeded networks at FDR < 0.05 were tested for overlap with PsychENCODE WGCNA modules, which capture neurobiological pathways and cell-types. Out of 34 gene-level (geneM) modules, for *C4A*-positive genes, the strongest enrichment was observed for astrocyte module (geneM3) at low copy number and for NFkB module (geneM5) at subsequently higher copy number. Microglial (geneM6) and interferon-response (geneM32) modules also showed stronger enrichment at higher copy number. Meanwhile for *C4A*-negative genes, we observed the strongest enrichment for synapse- and neuron-related modules. Text shows odds ratio from two-sided Fisher's exact test. Bonferroni-significant results are marked with black borders.

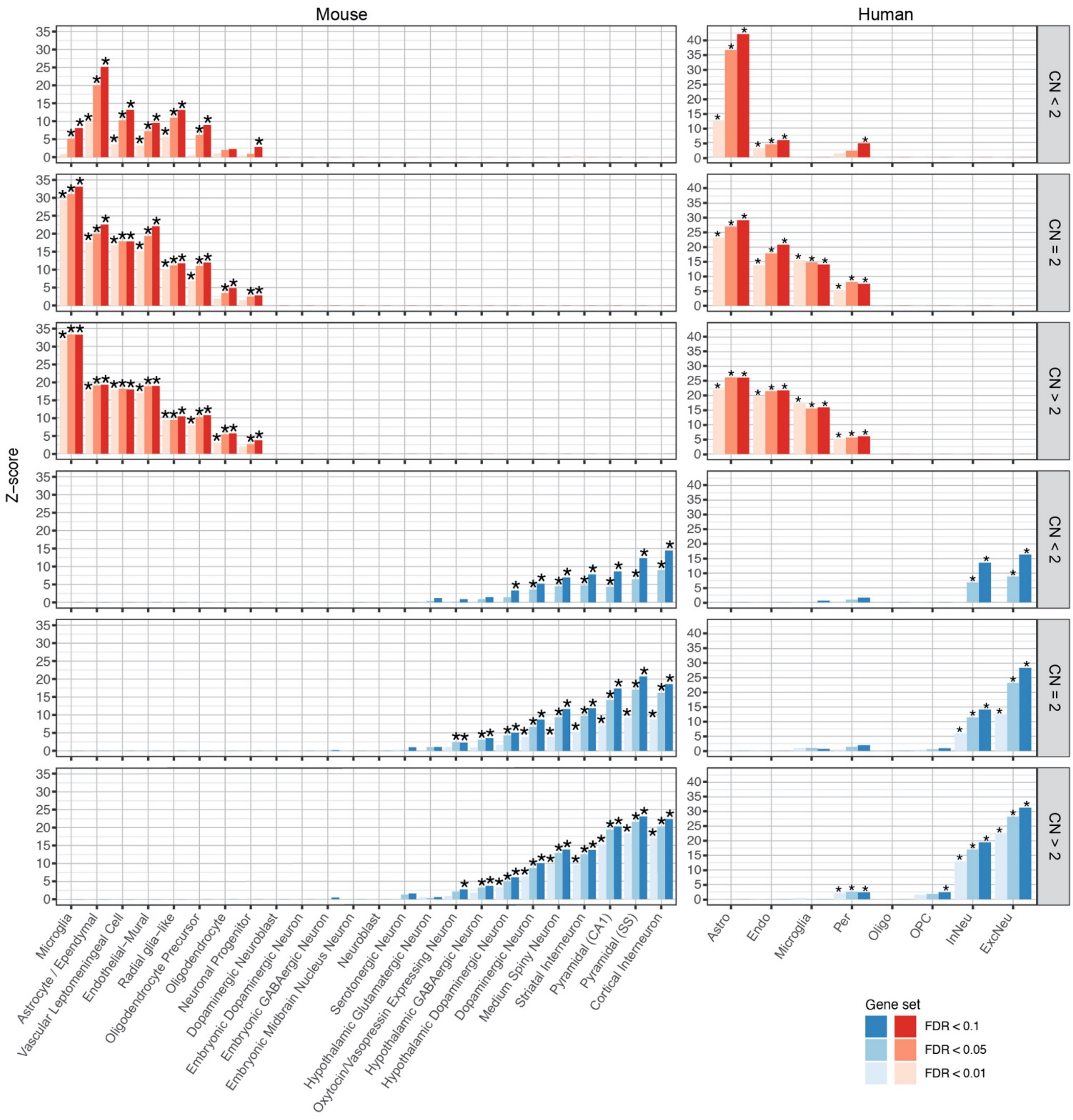


**Supplementary Figure 3. Expression of *C4A*-positive and *C4A*-negative genes in distinct mouse and human cell-types.** Gene sets obtained from the seeded networks were used for EWCE in mouse and human single-cell/nucleus RNA-seq data. All available major cell-types from either mouse or human brain were tested. Note that there were no cell-types from subcortical brain regions in the human dataset. To ensure that the observed enrichment pattern is preserved on a more global and systems-scale, gene sets obtained with more permissive FDR thresholds were also tested. *C4A*-positive and *C4A*-negative genes are shown in red and blue, respectively. Asterisks denote significance at FDR < 0.05.

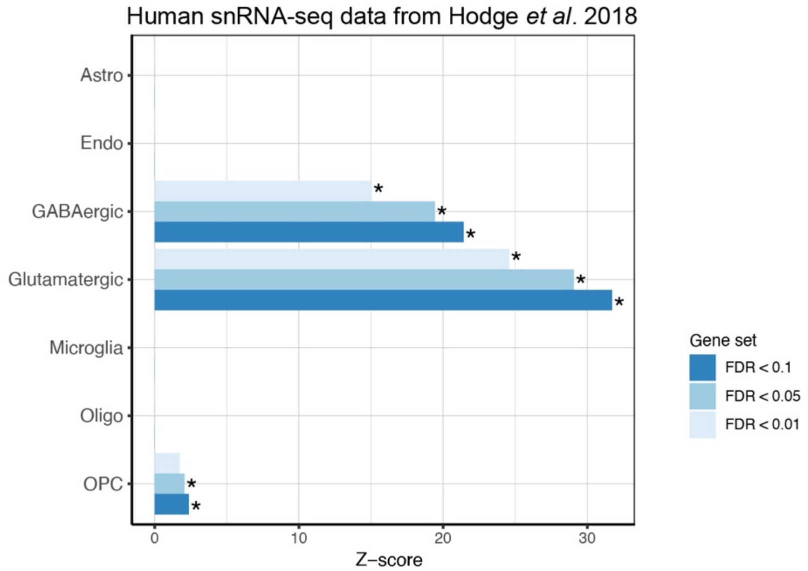


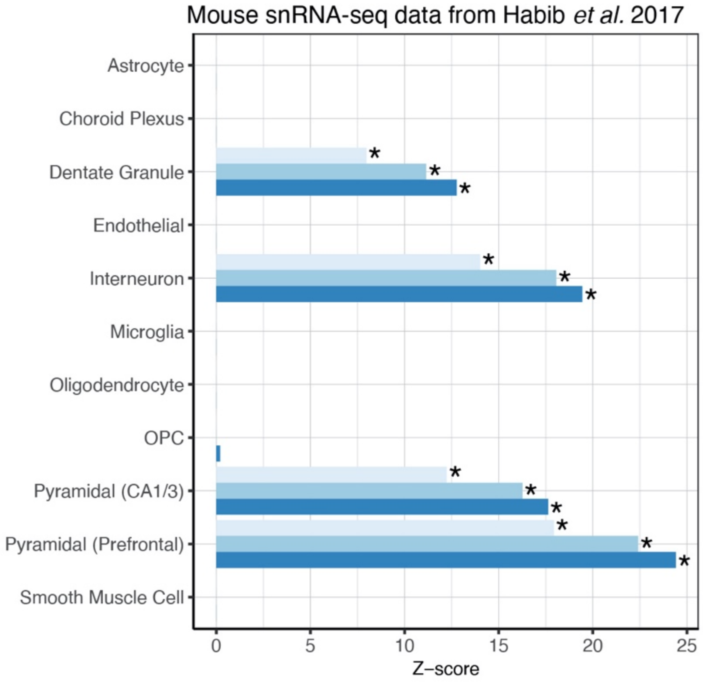


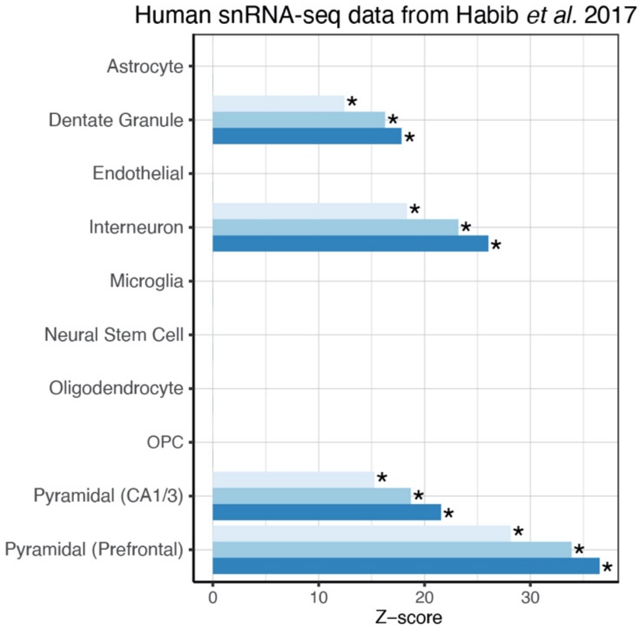


**Supplementary Figure 4. EWCE analyses in other single-cell/nucleus RNA-seq datasets.** *C4A*-negative genes obtained from high CNV group (i.e. CN > 2) at different FDR thresholds were used for EWCE in multiple single-cell/nucleus RNA-seq data. The results were consistent with previous analyses, where *C4A*-negative genes implicate neuronal and synaptic genes. Asterisks denote significance at FDR < 0.05.

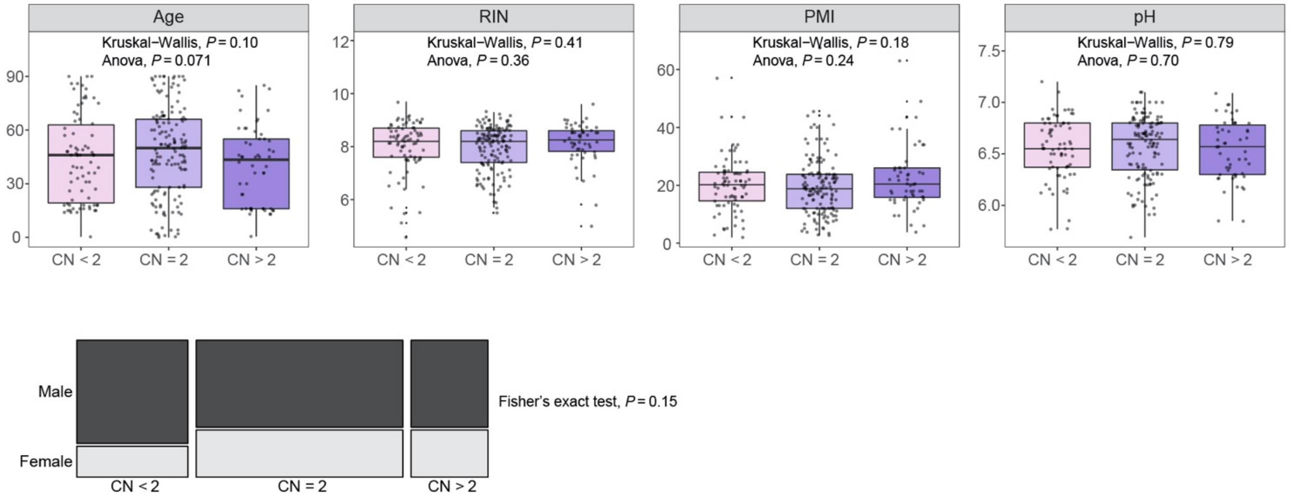


**Supplementary Figure 5. Baseline characteristics of control samples in PsychENCODE.** Age, RIN, postmortem interval (PMI), brain pH, and sex were balanced across the control samples (N = 78, 145, and 54 for CN < 2, CN = 2, and CN > 2, respectively). All boxplots show median and interquartile range (IQR) with whiskers denoting 1.5 × IQR.

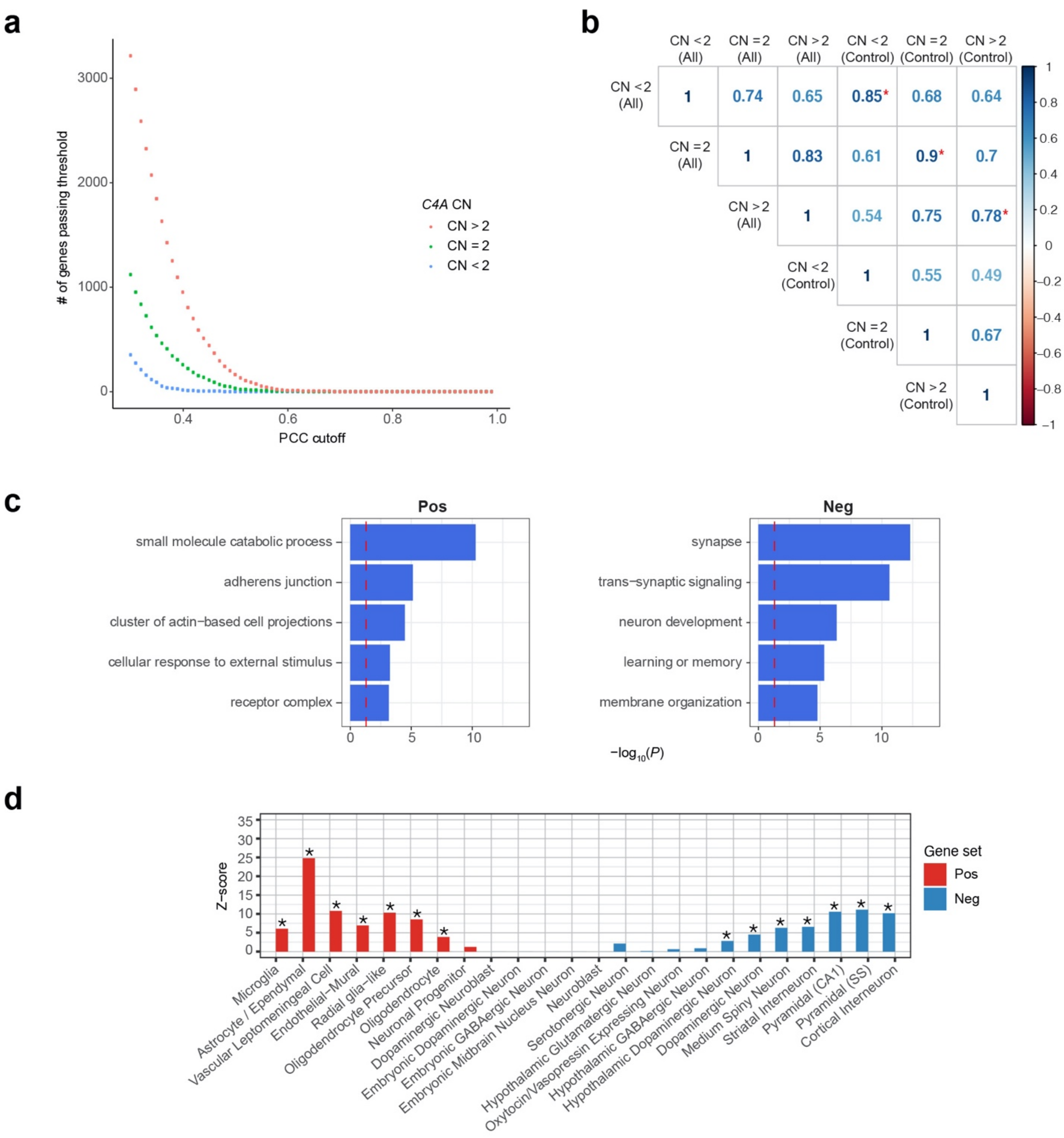


**Supplementary Figure 6**. **Characteristics of control-only *C4A*-seeded networks**. **a**, Networks constructed using only the control samples in PsychENCODE expanded in size irrespective of Pearson's correlation coefficient (PCC) threshold. **b**, Correlation of effect sizes (i.e. PCC) from all-sample (All) and control-only (Control) networks were the highest in pairs with equal copy numbers. **c**, Distinct pathways were enriched for *C4A*-positive and *C4A*-negative genes (top 500 genes each ranked by PCC) from control-only networks. **d**, Similarly, distinct cell-types were enriched for *C4A*-positive and *C4A*-negative genes (top 500) from control-only networks.
