## Extended Data Figures for "Brain gene co-expression networks link complement signaling with convergent synaptic pathology in schizophrenia"

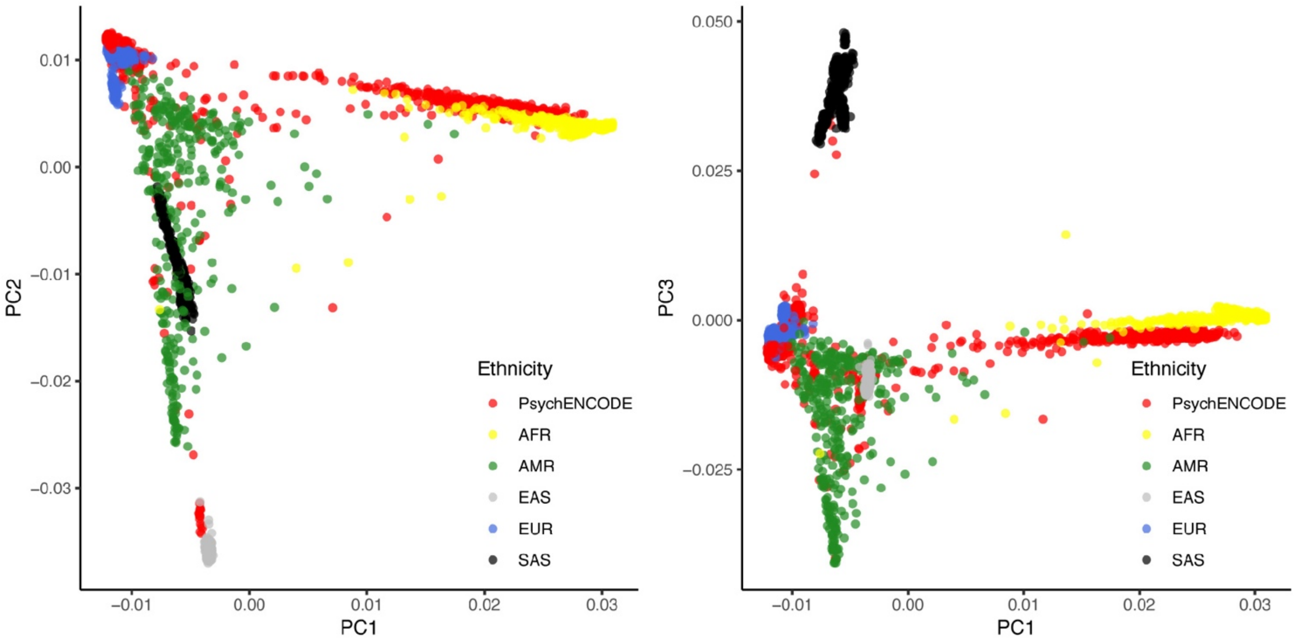


**Extended Data Fig. 1. Ancestry of PsychENCODE subjects.** Principal component analysis was performed using PLINK after merging the PsychENCODE genotype data with the 1000 Genomes Project reference panel. The PsychENCODE genotype data was available for a total 1,864 subjects to begin with. Each point represents an individual and points are color-coded by corresponding ethnicity. Global ancestry was inferred by k-nearest neighbors algorithm with the first five principal components. Downstream analyses were restricted to samples of European ancestry (N = 812).

| | CN < 2 | CN = 2 | CN > 2 |
|---|---|---|---|
| Autism (ASD) | 1 | 2 | 4 |
| Bipolar Disorder (BD) | 9 | 44 | 12 |
| Schizophrenia (SCZ) | 31 | 133 | 39 |
| Control (CTL) | 78 | 145 | 54 |
| Total | 119 | 324 | 109 |

**Extended Data Fig. 2. Number of PsychENCODE samples with high-quality *C4* imputation.** Total 552 samples had average imputed probabilistic dosage > 0.7. These samples were subsequently used to generate *C4A*-seeded networks.

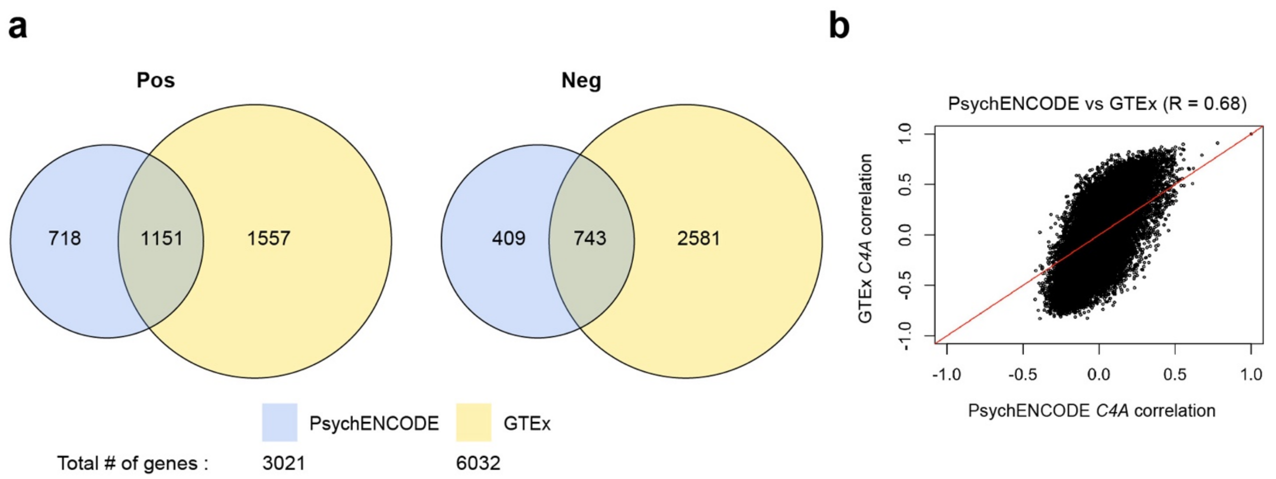


**Extended Data Fig. 3. Replication of PsychENCODE seeded network in GTEx. a**, Shown are Venn diagrams of the number of overlapping *C4A*-positive and *C4A*-negative genes in PsychENCODE and GTEx (OR's = 19 and 16, *P*'s < $10^{-16}$, respectively). These networks were constructed from frontal cortex samples of non-psychiatric controls with *C4A* CN = 2. **b**, Shown is correlation of effect sizes (i.e. PCC) of each gene that is shared between the two networks (R = 0.68, two-sided *P* < $10^{-16}$).

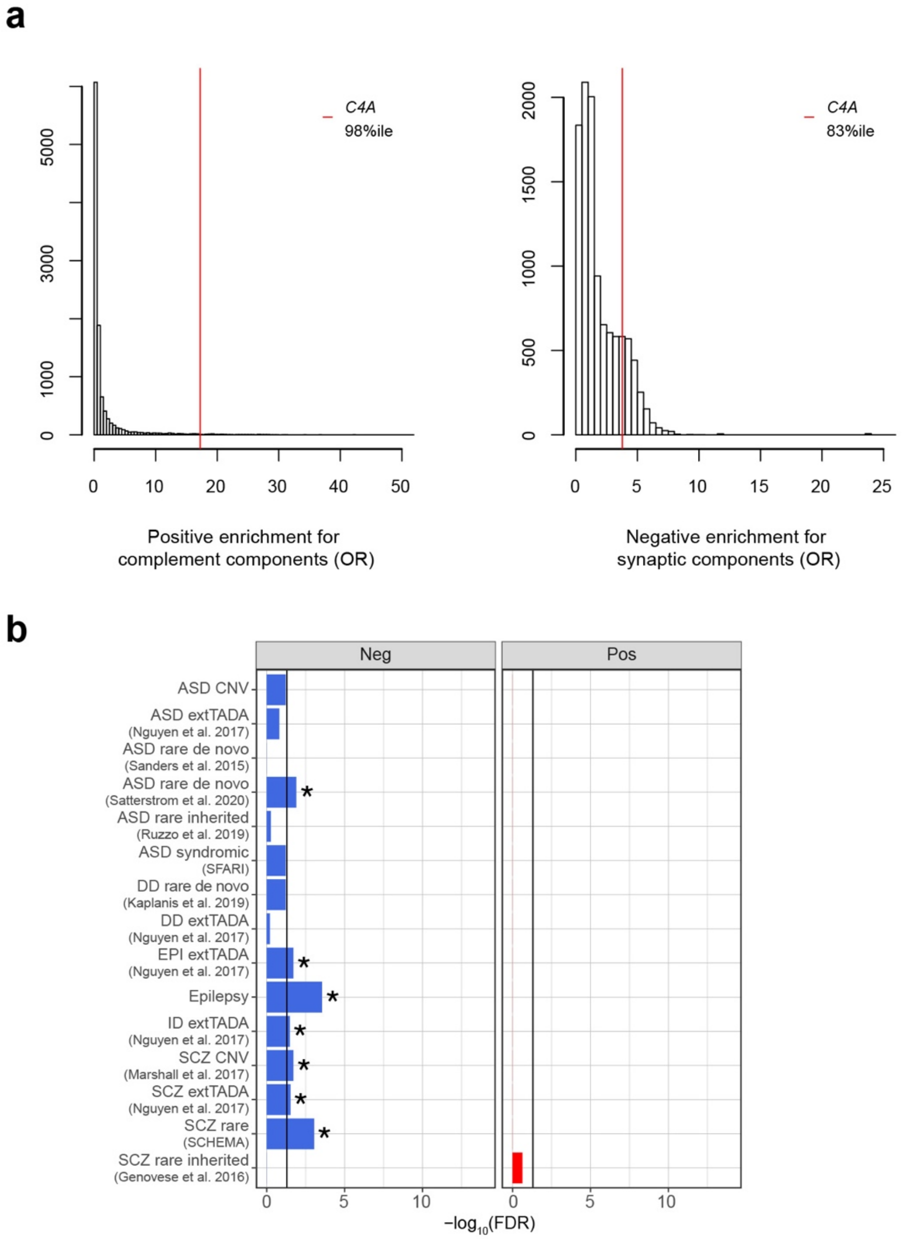


**Extended Data Fig. 4. Enrichment for complement components among *C4A*-positive genes and synaptic components as well as neurodevelopmental risk genes among *C4A*-negative genes. a**, Seed genes were permuted 10,000 times and corresponding seeded networks were tested for enrichment of the complement system (n = 57 genes) and synaptic components (n = 1,103 genes) from SynGo. Shown is distribution of the odds ratio from Fisher's exact test. **b**, *C4A*-positive and *C4A*-negative genes at FDR < 0.05 from the meta-analysis of PsychENCODE and GTEx were used for rare variant analyses (logistic regression with significance assessed through likelihood ratio test). The dotted line denotes FDR-adjusted *P* value at 0.05.

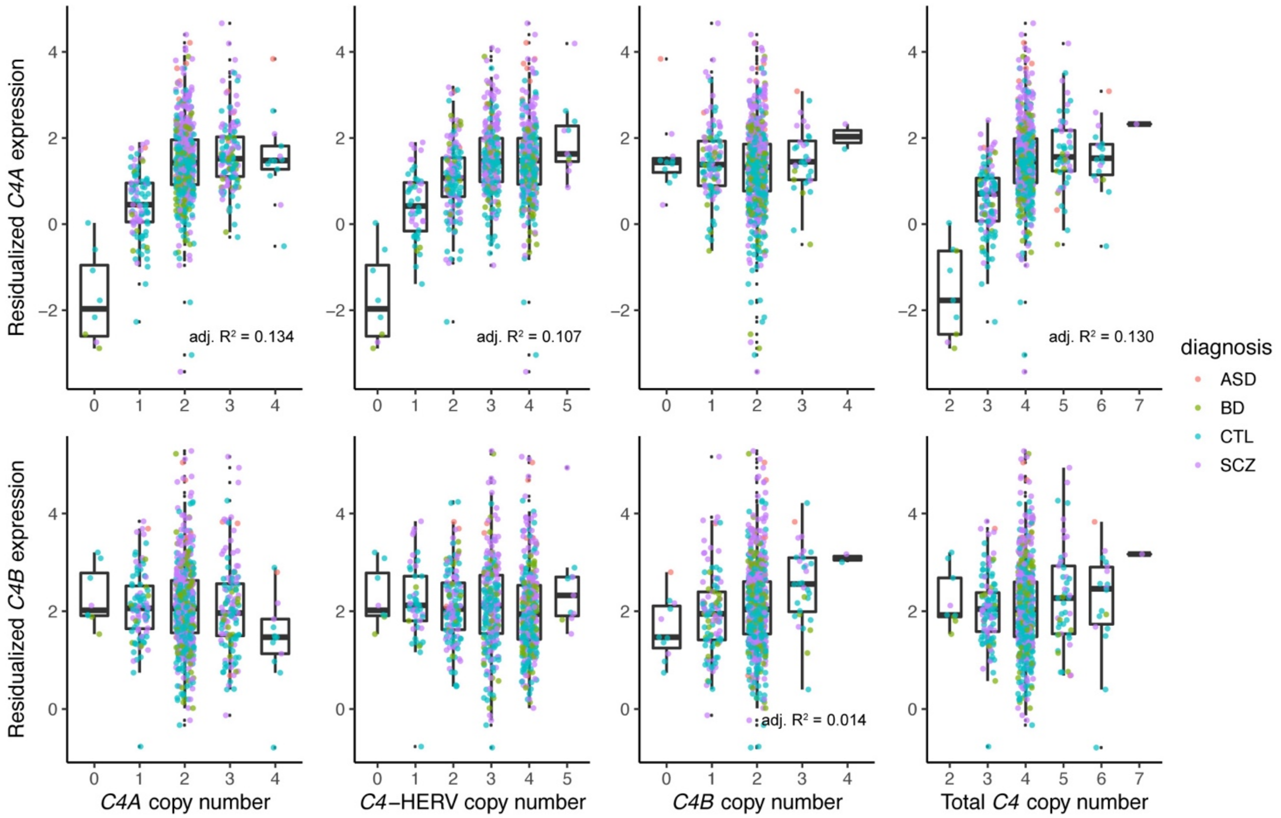


**Extended Data Fig. 5. Relationship between *C4* structural variation and *C4* gene expression.** Residualized *C4* gene expression (i.e. normalized and corrected for all known biological and technical covariates except the diagnosis status) was associated strongly with corresponding gene copy number (total N = 812; N = 20, 114, 367, and 311 for ASD, BD, CTL, and SCZ samples, respectively). Adjusted $R^2$ values are shown for significant correlations. Of note, the best linear models for *C4A* and *C4B* expression explained up to 22% and 2.7% of variation in expression, respectively. All boxplots show median and interquartile range (IQR) with whiskers denoting 1.5 × IQR.

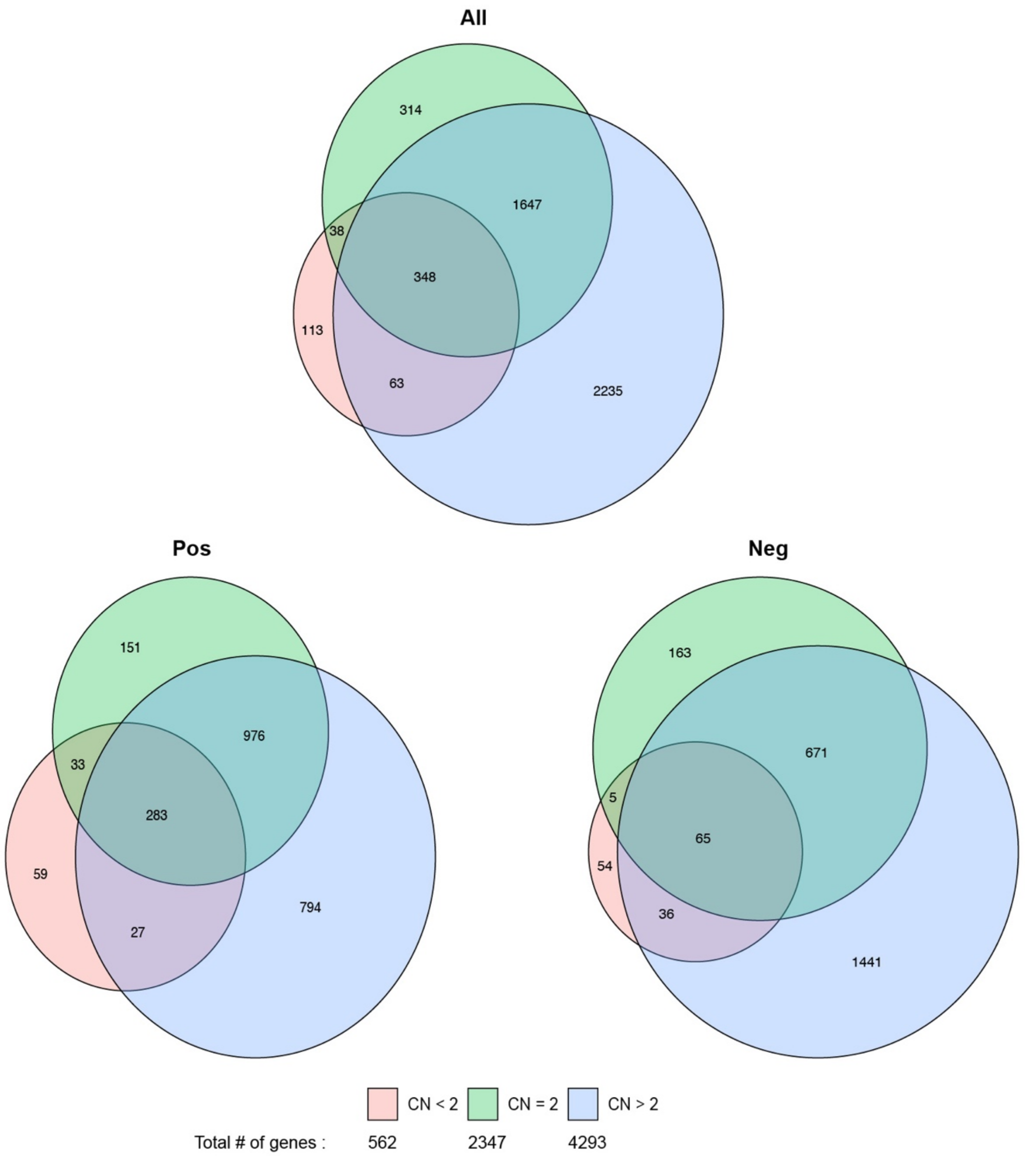


**Extended Data Fig. 6. Larger number of *C4A*-positive and *C4A*-negative genes with increased *C4A* copy number**. Shown are Venn diagrams of the number of overlapping *C4A*-positive and *C4A*-negative genes across three CNV groups. Note that the sum of positive and negative genes is equal to the total number of co-expressed genes. The size of the circle is approximately proportional to the number of genes.

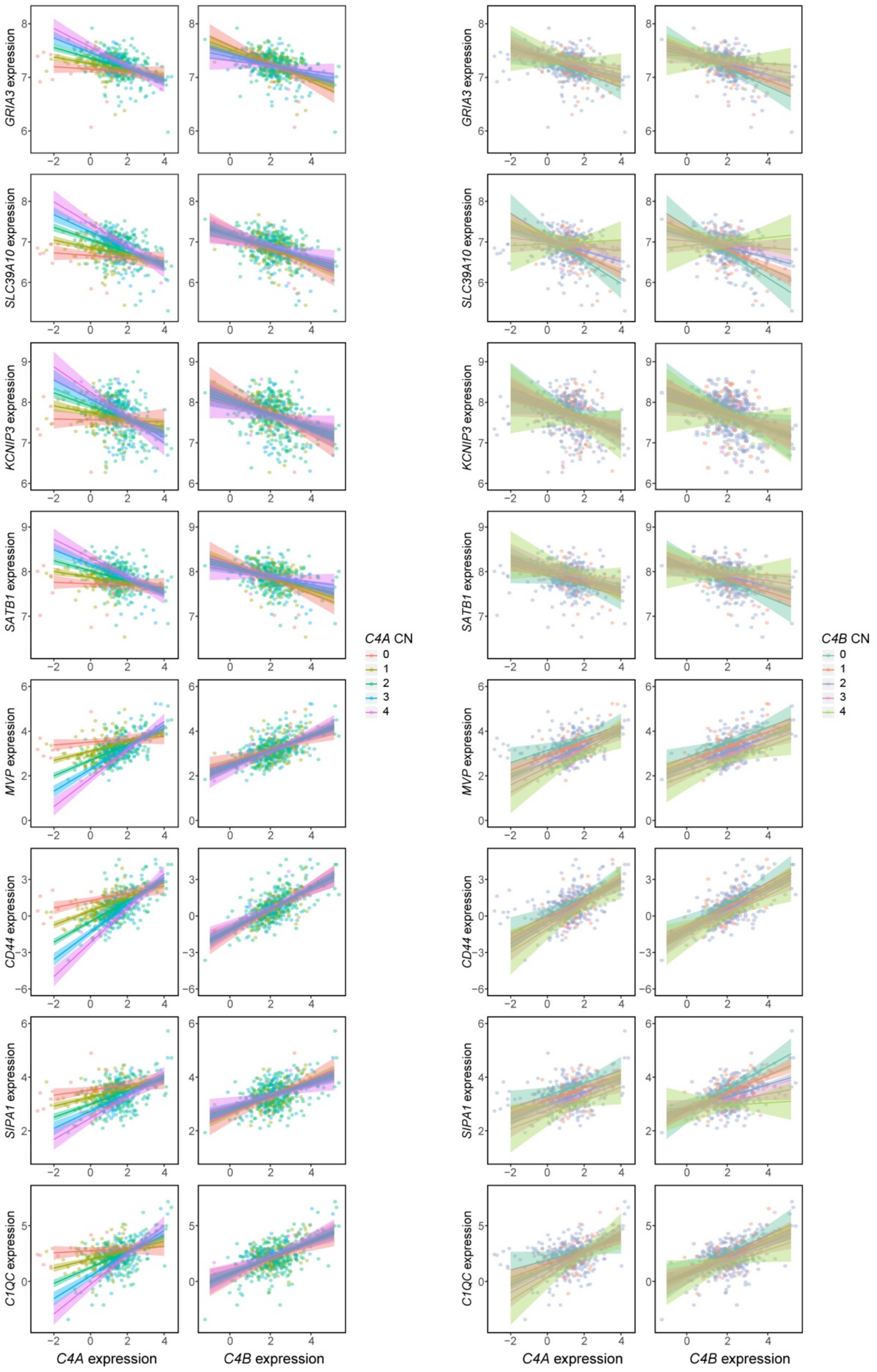


**Extended Data Fig. 7. *C4A*-specific interaction with *C4A* copy number.** Multiple regression was performed with interaction terms between *C4* copy numbers and *C4* gene expression. Significant interaction effect was present only between *C4A* copy number and *C4A* expression. Several genes are highlighted to demonstrate this interaction. Also shown are fitted linear models with 95% confidence bands.

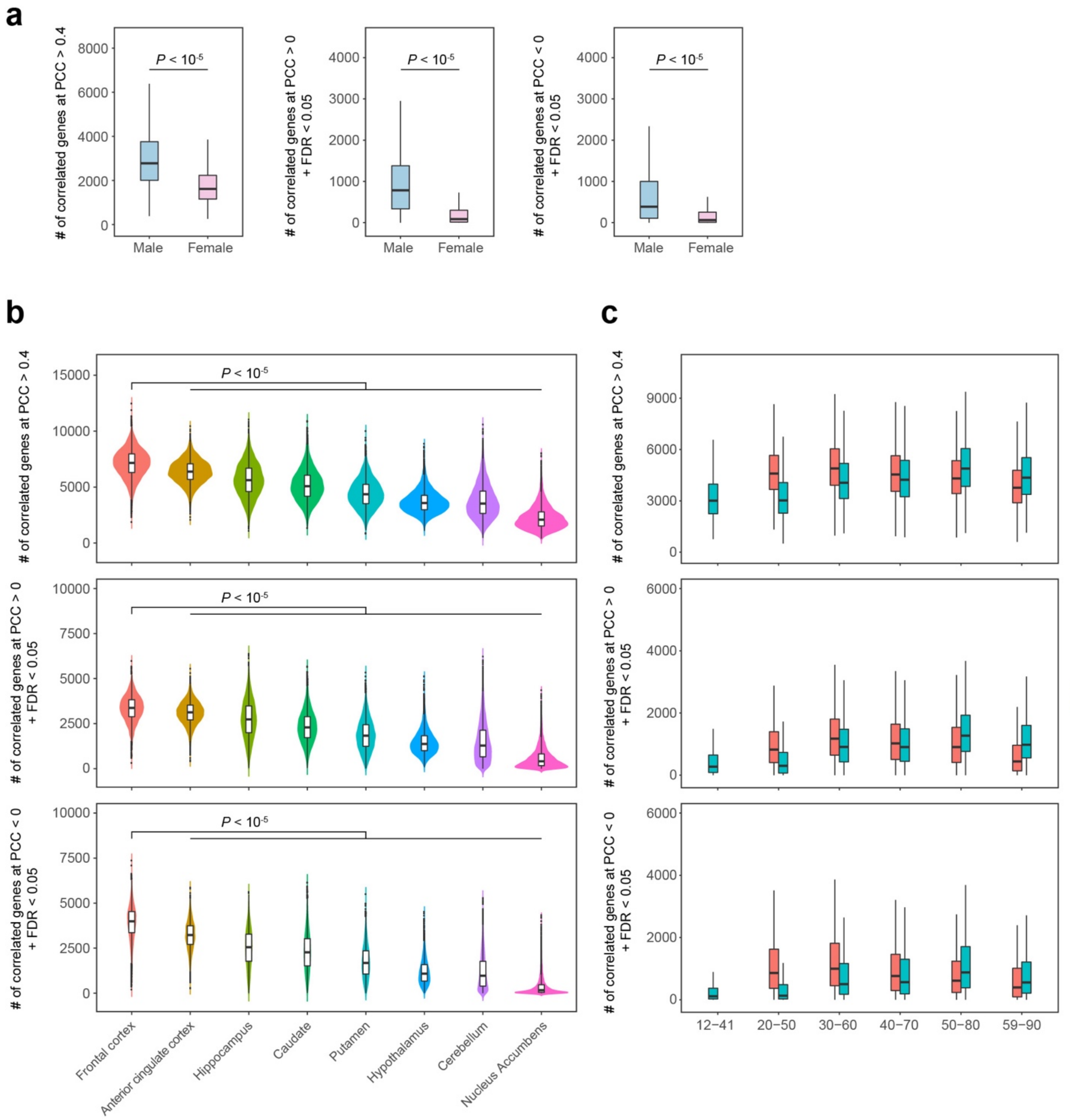


**Extended Data Fig. 8. Sex and spatiotemporal differences in *C4A* co-expression. a**, Three different thresholds were tested, namely the number of total co-expressed genes at PCC > 0.4 and the number of *C4A*-positive and *C4A*-negative genes at FDR < 0.05. Males had more co-expressed genes than females regardless of the threshold metric used (N = 36, 38, 45, 47, 39, 45, 39, and 45 for frontal cortex, anterior cingulate cortex, hippocampus, caudate, putamen, cerebellum, hypothalamus, and nucleus accumbens, respectively; permutation test, $P < 10^{-5}$). **b**, Similarly, frontal and anterior cingulate cortex were the two most connected regions for *C4A* regardless of the threshold metric used (N = 36, 38, 45, 47, 39, 45, 39, and 45 for frontal cortex, anterior cingulate cortex, hippocampus, caudate, putamen, cerebellum, hypothalamus, and nucleus accumbens, respectively; permutation test, $P < 10^{-5}$). **c**, Leftward shift in co-expression peak was observed in SCZ cases compared to neurotypical controls across different threshold metrics (N = 30, 42, 57, 68, 47, and 32 for control samples in each age bin; N = 36, 46, 55, 45, and 47 for SCZ samples). All boxplots show median and interquartile range (IQR) with whiskers denoting 1.5 × IQR.

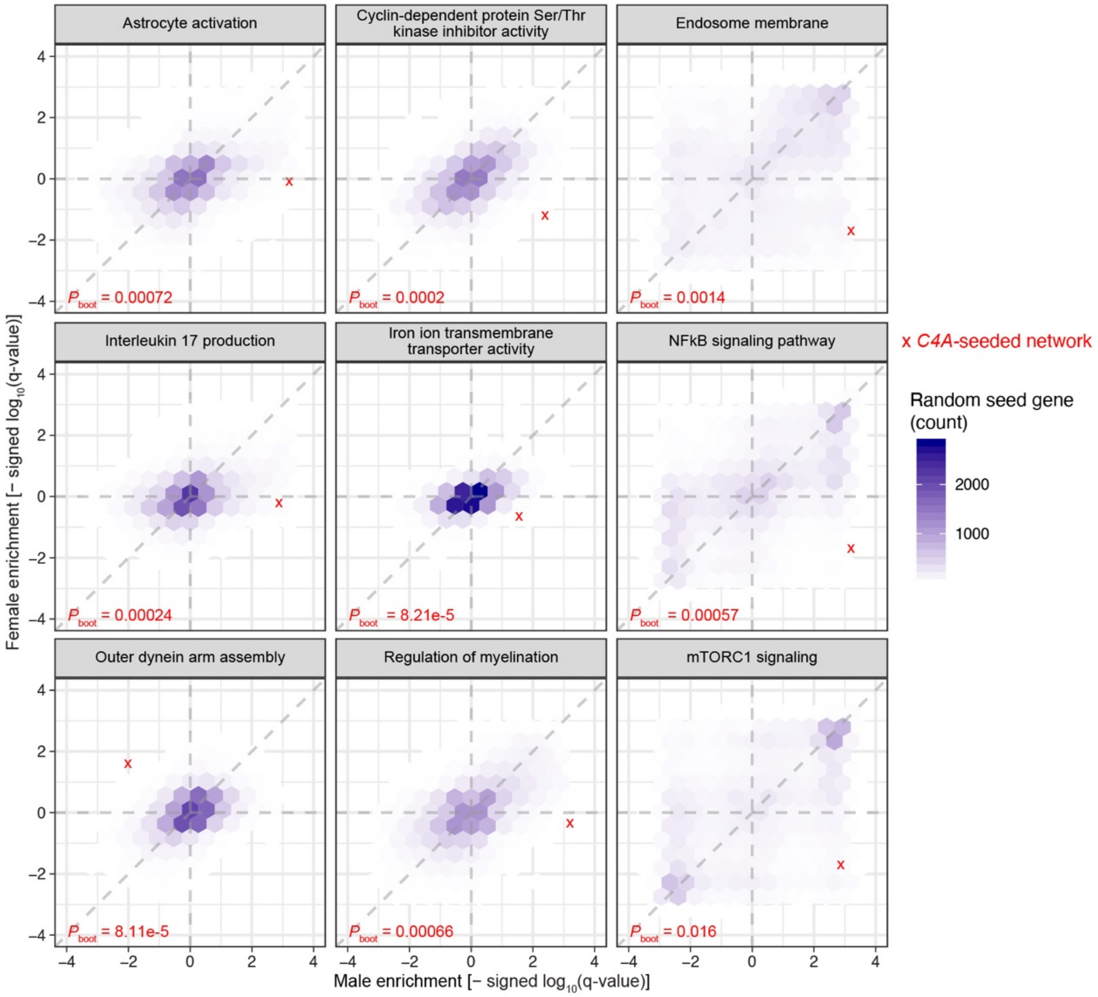


**Extended Data Fig. 9. Pathways exhibiting differential co-expression in males and females.** Shown are GSEA enrichments for *C4A* compared to 10,000 random seed genes. Genes were ranked by the magnitude of co-expression in male and female networks separately, and the corresponding gene list was used for GSEA. Several pathways showed the opposite direction of effect.

| | BD (All) | SCZ subsample (BD matched) | SCZ (All) | SCZ (EUR) | SCZ (All) \| *C4A* expr | SCZ (EUR) \| *C4A* expr | SCZ (EUR) \| *C4A* CN | SCZ (EUR) \| *C4A* CN + *C4A* expr |
|---|---|---|---|---|---|---|---|---|
| C1QA | −0.14 | −0.0037 | 0.013 | 0.049 | −0.11 | −0.08 | 0.041 | −0.08 |
| C1QB | −0.043 | 0.05 | 0.074 | 0.1 | −0.072 | −0.044 | 0.099 | −0.044 |
| C1QC | 0.015 | 0.054 | 0.073 | 0.087 | −0.075 | −0.065 | 0.081 | −0.065 |
| C1R | 0.1 | 0.25 * | 0.25 * | 0.23 * | 0.13 * | 0.12 * | 0.23 * | 0.12 * |
| C1S | 0.039 | 0.13 * | 0.13 * | 0.11 * | 0.059 * | 0.039 | 0.11 * | 0.038 |
| C2 | 0.061 | 0.042 | 0.044 | 0.025 | −0.021 | −0.038 | 0.024 | −0.039 |
| C3 | −0.18 | −0.21 * | −0.19 * | −0.18 * | −0.29 * | −0.29 * | −0.19 * | −0.29 * |
| C3AR1 | −0.18 | −0.2 * | −0.18 * | −0.17 | −0.26 * | −0.25 * | −0.18 | −0.25 * |
| C4A | 0.071 | 0.4 * | 0.39 * | 0.44 * | 0 | 0 | 0.37 * | 0 |
| C4B | 0.064 | 0.34 * | 0.33 * | 0.34 * | 0.095 * | 0.1 | 0.35 * | 0.1 |
| C5 | 0.0061 | −0.03 | −0.023 | −0.03 | −0.031 | −0.039 | −0.03 | −0.039 |
| C5AR1 | 0.029 | −0.00054 | 0.02 | 0.072 | −0.061 | −0.0087 | 0.068 | −0.0092 |
| C5AR2 | −0.079 | −0.15 * | −0.13 * | −0.11 | −0.15 * | −0.14 * | −0.11 | −0.14 * |
| C7 | 0.037 | 0.11 | 0.094 | 0.018 | 0.01 | −0.07 | 0.013 | −0.071 |
| C8G | −0.093 | −0.078 | −0.068 | 0.042 | −0.052 | 0.065 | 0.041 | 0.066 |
| CD46 | 0.012 | −0.024 | −0.024 | −0.031 | −0.021 | −0.027 | −0.031 | −0.027 |
| CD55 | −0.0039 | −0.018 | −0.029 | −0.013 | −0.046 | −0.026 | −0.016 | −0.026 |
| CD59 | 0.027 | 0.054 * | 0.055 * | 0.053 * | 0.032 | 0.03 | 0.054 * | 0.029 |
| CD93 | 0.092 | 0.092 | 0.11 | 0.085 | −0.024 | −0.038 | 0.08 | −0.04 |
| CFB | −0.094 | 0.047 | 0.06 | 0.03 | −0.023 | −0.039 | 0.027 | −0.04 |
| CFD | −0.17 * | −0.071 | −0.077 | −0.057 | −0.094 | −0.081 | −0.055 | −0.081 |
| CFH | −0.13 | 0.034 | 0.035 | −0.027 | −0.024 | −0.093 | −0.028 | −0.094 |
| CFI | −0.087 | 0.087 | 0.091 | 0.079 | −0.042 | −0.05 | 0.077 | −0.053 |
| CFP | −0.04 | −0.0095 | −0.0024 | −0.019 | 0.011 | 0.0011 | −0.0098 | 0.00053 |
| CLU | 0.088 * | 0.11 * | 0.12 * | 0.12 * | 0.072 * | 0.072 * | 0.12 * | 0.071 * |
| COLEC10 | 0.033 | −0.13 | −0.11 * | −0.13 | −0.09 | −0.12 | −0.14 | −0.12 |
| COLEC11 | 0.019 | 0.054 | 0.058 | 0.21 | 0.053 | 0.19 | 0.21 | 0.18 |
| CR1 | −0.17 | 0.19 * | 0.14 * | 0.26 * | 0.022 | 0.14 | 0.26 * | 0.14 |
| CSMD1 | 0.0037 | 0.0028 | 0.00099 | −0.02 | 0.014 | −0.01 | −0.019 | −0.01 |
| CSMD2 | 0.038 | −0.0019 | −0.003 | −0.0045 | 0.0035 | 0.0053 | −0.0022 | 0.0052 |
| CSMD3 | 0.0044 | 0.00025 | 0.0026 | 0.0025 | 0.012 | 0.013 | 0.0055 | 0.013 |
| ELANE | −0.045 | −0.13 | −0.089 | −0.1 | −0.11 | −0.11 | −0.094 | −0.11 |
| FCN1 | 0.017 | 0.15 | 0.16 | 0.34 * | 0.071 | 0.25 | 0.34 * | 0.25 |
| FCN3 | −0.055 | 0.067 | 0.066 | 0.031 | 0.0041 | −0.04 | 0.023 | −0.041 |
| ITGAM | −0.14 | −0.19 * | −0.18 * | −0.19 * | −0.23 * | −0.25 * | −0.19 * | −0.25 * |
| ITGAX | −0.14 | −0.33 * | −0.31 * | −0.3 * | −0.32 * | −0.32 * | −0.3 * | −0.32 * |
| ITGB2 | −0.15 | −0.14 | −0.12 | −0.14 | −0.22 * | −0.24 * | −0.14 | −0.24 * |
| MASP1 | 0.12 * | 0.095 | 0.11 * | 0.049 | 0.028 | −0.032 | 0.052 | −0.033 |
| MASP2 | 0.011 | −0.017 | −0.016 | −0.02 | −0.023 | −0.045 | −0.024 | −0.045 |
| SERPING1 | 0.13 * | 0.16 * | 0.16 * | 0.13 * | 0.098 * | 0.072 | 0.13 * | 0.071 |
| VSIG4 | 0.008 | 0.11 | 0.13 | 0.17 | −0.0089 | 0.03 | 0.16 | 0.03 |
| VTN | 0.054 | −0.048 | −0.048 | −0.042 | −0.03 | −0.026 | −0.038 | −0.026 |

$-\log_{10}$FDR: 0.0 2.5 5.0 7.5

**Extended Data Fig. 10. Differential gene expression of the complement system in SCZ and BD**. Differential expression (DE) for brain-expressed complement system genes (n = 42) was assessed in SCZ (N = 531) and BD (N = 217) compared to controls (N = 895). DE was repeated for SCZ after randomly downsampling to match the sample size of BD. DE was also repeated for SCZ while adjusting for *C4A* expression and/or *C4A* copy number. Since *C4A* copy number was only imputed for samples of European ancestry, a subset of PsychENCODE samples was used for such conditional analyses (N = 311 and 367 for SCZ and controls, respectively). Text shows $\log_2$FC. Asterisks denote significance at FDR < 0.1.
